# Continuous foraging behavior shapes patch-leaving decisions in pigeons: A 3D tracking study

**DOI:** 10.64898/2026.02.18.706261

**Authors:** Guillermo Hidalgo-Gadea, Onur Güntürkün, Mary Flaim, Patrick Anselme

**Affiliations:** Department of Biopsychology, Faculty of Psychology, Ruhr University Bochum, Bochum, Germany

**Author notes:** Correspondence concerning this article should be addressed to Guillermo Hidalgo Gadea, Faculty of Psychology, Ruhr University Bochum, Universitätsstraße 150, 44801 Bochum, Germany. These authors contributed equally to this study and share senior authorship (Mary Flaim, Patrick Anselme).

**Keywords:** optimal foraging, marginal value theorem, travel cost, pigeon behavior, 3D pose tracking, cognition

## Abstract

Optimal foraging behavior is a key component of successful adaptations to natural environments. Understanding how animals decide to stay near food or to leave it for another food patch gives us insights into the underlying cognitive mechanisms that govern adaptive behaviors. 3D pose tracking was used to determine how pigeons exploit a 4 square meter arena with two separate platforms (i.e. food patches) whose absolute and relative elevations were manipulated. Detailed kinematic features of foraging and traveling behaviors were quantified using automated video tracking, without a need for manual coding. Our computational approach captured continuous, high-dimensional movement patterns and enabled precise quantification of travel costs between patches. Combined with mixed-effects survival analysis, our fine-grained behavioral tracking provided detailed insight into the moment-by-moment dynamics of patch-leaving decisions of pigeons. As expected from behavior optimization models, our results showed a preference to visit a ground food platform first, and longer latencies to leave an elevated platform. Foraging activity significantly decreased throughout the session, with shorter visits, less pecks per visit, and a decrease in inter-peck variability. However, a mixed-effects Cox regression modeled pigeons’ patch-leaving probability, demonstrating that current and cumulative foraging parameters between patches significantly enhanced the model’s predictive power beyond patch accessibility (i.e., beyond travel costs). This suggests that pigeons integrate both current environmental cues and their individual foraging history when making patch-leaving decisions. Our findings are discussed in relation to the marginal value theorem and optimal foraging theory.

---

Food distribution in the environment is more often patchy than homogeneous, meaning that finding a food item in one location increases the likelihood of finding other food items nearby. To maximize profitability of local search behavior, animals have evolved decision rules that help them optimize prey selection (e.g., Richardson & Verbeek, 1986; Meire & Ervynck, 1986) or adjust their risk-taking behavior based on energy reserves and other factors (e.g., Barnard & Brown, 1985; Caraco et al., 1990; Cartar & Dill, 1990). Some foraging heuristics include leaving a patch after consuming a fixed number of food items (Gibb, 1962), after a fixed duration (Krebs, 1973), or after a period without food encounters (Krebs et al., 1974). Importantly, such decision rules can emerge dynamically from interactions between an organism and its environment rather than necessarily relying on internalized representations of the foraging habitat. For example, the slime mold (*Physarum polycephalum*), a giant amoeba, exhibits optimal foraging behavior using simple heuristics based solely on information obtained through foraging itself (Latty & Beekman, 2015). This macroscopic unicellular organism can solve shortest path problems (Nakagaki et al., 2000) and balance its uptake of multiple macronutrients (Dussutour et al., 2010) without a single neuron. While such examples illustrate that complex foraging patterns can emerge from simple rules, in vertebrates these behaviors are typically embedded within flexible decision-making systems that integrate past experience and current environmental information.

Another well-studied decision rule in animal foraging is determining the optimal time to stay in a food patch before moving to another, given the foraging costs—such as time, effort, and risk—that this move may represent. The marginal value theorem (MVT) estimates optimal residence time in a patch, given the distance to the next patch (Charnov, 1976). The MVT states that a patch should be left when the marginal gain (the rate of increase in the food amount obtained per unit time) in that patch drops below the average gain that could be obtained by moving to a new patch. By balancing the benefits of exploiting a patch with the costs of staying there too long, the individual can optimize its foraging efficiency and maximize its net energy intake. For that, the animal needs to estimate the quality and quantity of resources in other potential patches and the time it would take to reach them. This estimation is based on prior knowledge of the environment.

In theory, the MVT does not assume that an animal has omniscient knowledge of its environment; rather, foraging decisions are based on the information available at a given time, even if that information is limited and imperfect. When an animal visits multiple patches during a single foraging bout, it can form an estimate of the environment based on the proportion of food items found per unit time in the visited patches. This past information becomes useful for future decisions (MacArthur, 1972). In practice, however, in the absence of a well-controlled environment, estimating what an animal knows or does not know is rather difficult. As a result, any deviation from the MVT’s predictions might be attributed to insufficient background knowledge about the environment instead of being a failure of the theory—and vice-versa. Thus, rigorous tests of the MVT require experimental designs that constrain uncertainty and allow clearer inference about the information available to the animal.

The matching law (Herrnstein, 1961; Baum, 1974), which posits that behavior rates between several options (“patches”) should be proportional to reinforcement rates provided by those options, faces a similar challenge. Various constraints—both identified and unidentified—arising from an animal’s estimates can lead to deviations from theoretical predictions (e.g., Anselme et al., 2022; McDowell, 2013; Strand et al., 2022). To minimize these deviations, experimental designs should carefully control food availability and effort requirements while ensuring that the foraging context remains ecologically relevant. For example, pecking at spatially distinct food locations more closely resembles natural foraging than laboratory tasks that require withholding responses to delayed reinforcement.

If foraging behavior is aligned with the matching law (Dallery & Baum, 1991), activity levels should be proportional to reinforcement rates and should exhibit extinction-like patterns when reinforcement decreases—such as shorter visit durations in depleted patches. However, deviations from strict matching have been observed in semi-naturalistic environments—at least in the short term. For example, pigeons have been shown to display overmatching, foraging more than expected in a patch whose caches were inconsistently baited with food compared to a patch in which each cache contained a food item (Anselme et al., 2022, 2024). There is also evidence that a reduction in food probability favors search behavior, notably through an increase in behavioral variability and vigor (Anselme et al., 2013; Blaisdell et al., 2016; Bond, 1983; Wittek et al., 2022).

Recent advances in automated behavioral tracking, including multi-camera motion capture and markerless pose-estimation approaches such as DeepLabCut (Mathis et al., 2018; Nath et al., 2019) and related frameworks (Chimento et al., 2025; Delacoux & Kano, 2024; Karashchuk et al., 2021; Naik et al., 2023), have enabled fine-grained quantification of continuous behavioral dynamics without invasive equipment (see also Hidalgo-Gadea et al., (2026) for a short review). Building on these developments, we adapted our own automated multi-camera 3D tracking system to quantify pigeons’ pose, movement and pecking dynamics during a foraging task. This non-invasive approach allowed us to link moment-to-moment foraging activity to patch-leaving decisions and to examine how transition costs and cumulative foraging history shape decision structure.

In this study, we investigated the foraging behavior of pigeons (*Columba livia*) in a well-controlled environment designed to approximate semi-naturalistic conditions. We tested several predictions from Charnov’s (1976) MVT using continuous 3D tracking to quantify both movement and foraging activity, to identify how pigeons integrate current patch information and cumulative experience when making decisions. The task consisted of two food patches (30 baited holes per platform) that were arranged either both on the ground (0–0), both elevated at 75 cm (75–75), or asymmetrically (0–75). We manipulated travel costs between two food patches by varying their relative elevation, allowing us to examine how accessibility influences patch-leaving behavior. Allowing repeated revisits within a 20-minute session further extends traditional MVT paradigms that often restrict patch access (Hong & Wolfe, 2024; but see Bracis et al., 2018).

Based on optimal foraging theory and the assumption that pigeons dynamically evaluate travel costs and diminishing returns, we derived the following predictions. First, pigeons engage in spontaneous foraging behavior and favor the platform associated with minimal travel costs. This suggests that the ground platform should be visited before the elevated platform in the asymmetric elevation condition (0-75), while no preference should be observed between the platforms in the symmetric conditions (0-0 and 75-75). Second, foraging activity differs based on the accessibility of a food patch and the travel cost between platforms. In particular, the area covered on a ground platform should be larger if the other platform is elevated rather than on the ground (because of higher transit costs), and the time spent on a platform before leaving it should be shorter in condition 0-0 than in the other two conditions (because of lower transit costs in the absence of flight) and shorter in condition 75-75 than in condition 0-75 (because horizontal flight is less costly than vertical flight). Furthermore, platforms whose access requires less effort facilitate foraging activity, and pigeons should eat more food items on ground platforms than on elevated platforms. Similarly, pigeons should transit more often in conditions 0-0 and 75-75 than in condition 0-75 because of lower transit costs. Third, foraging activity decreases as the platforms become food depleted towards the end of a session. In other words, the time spent, and the distance traveled on any platform should decrease over time. However, pecking activity should become more variable and pecking intensity per unit time should increase over a session. The pigeons should also be more often outside of a platform later in a session, switching from foraging to exploratory behavior when patches have been depleted. Fourth, we tested whether current foraging activity (as a proxy for intake rate), platform condition (as a proxy for transit cost), and cumulative foraging history (further referred to as overpecking) jointly predict patch-leaving decisions. Thus, this would provide a useful estimate of the current and cumulative rate of food intake in semi-naturalistic environments based on pigeons’ foraging investment (pecking activity, number of visits, and time spent).

## Method

### Animals and housing conditions

Twelve adult homing pigeons (10 females; age: 6.2±4.1 years) obtained from local breeders, were maintained at 85-90% of their free-feeding weight (i.e. experimental weight) for the duration of the experiment. Water was accessible ad libitum in their home cage. Two pigeons were housed individually, while the other 10 individuals were housed in two separate group aviaries, all under a 12h light/dark cycle (lights on at 7:30 am). The pigeons were used in a previous experiment involving the same foraging platforms, but where no choices had to be made, ensuring that the present task involved novel decision demands. The experiments were carried out in compliance with the European Communities Council Directive of September, 22 2010 (2010/63/EU) and the specifications of the German law for the prevention of cruelty to animals. They were also approved by the animal ethics committee of the Landesamt für Natur, Umwelt und Verbraucherschutz (LANUV) NRW, Germany. We confirm that all methods were carried out in accordance with relevant guidelines and regulations and that the study was conducted in compliance with the ARRIVE guidelines.

### Apparatus

We used a 2 meter wide and 2 meter high hexagonally shaped arena fitted with two wooden foraging platforms (60 cm x 50 cm) (Figure 1). Each platform had 30 symmetrically distributed holes, 1.5 cm wide and 1.5 cm deep, each containing a single food item—for a total 60 mixed green and yellow peas (approx. 10 grams, 15kcal, 24% carbohydrate, 0.7% fat, 10% protein). The platforms were covered with black adhesive film and the holes were cut open with x-shaped slits, leaving the food items accessible through flaps of the occlusive cover but not visible from a distance— as described in Anselme et al. (2022). Both foraging platforms were positioned at opposite walls of the area, either on the ground or on 75 cm high aluminum legs. One platform was always located to the left of a Murphy door, hidden on one of the walls. All six walls were visually indistinguishable from each other and fully enclosed with a solid white ceiling panel. Six synchronized RGB cameras (Teledyne FLIR, BFS-U3-16S2C-CS) were mounted around the ceiling of the arena, one on each wall, and recorded at 50 Hz and 1440:1080px resolution throughout the experiment.

**Figure 1.**
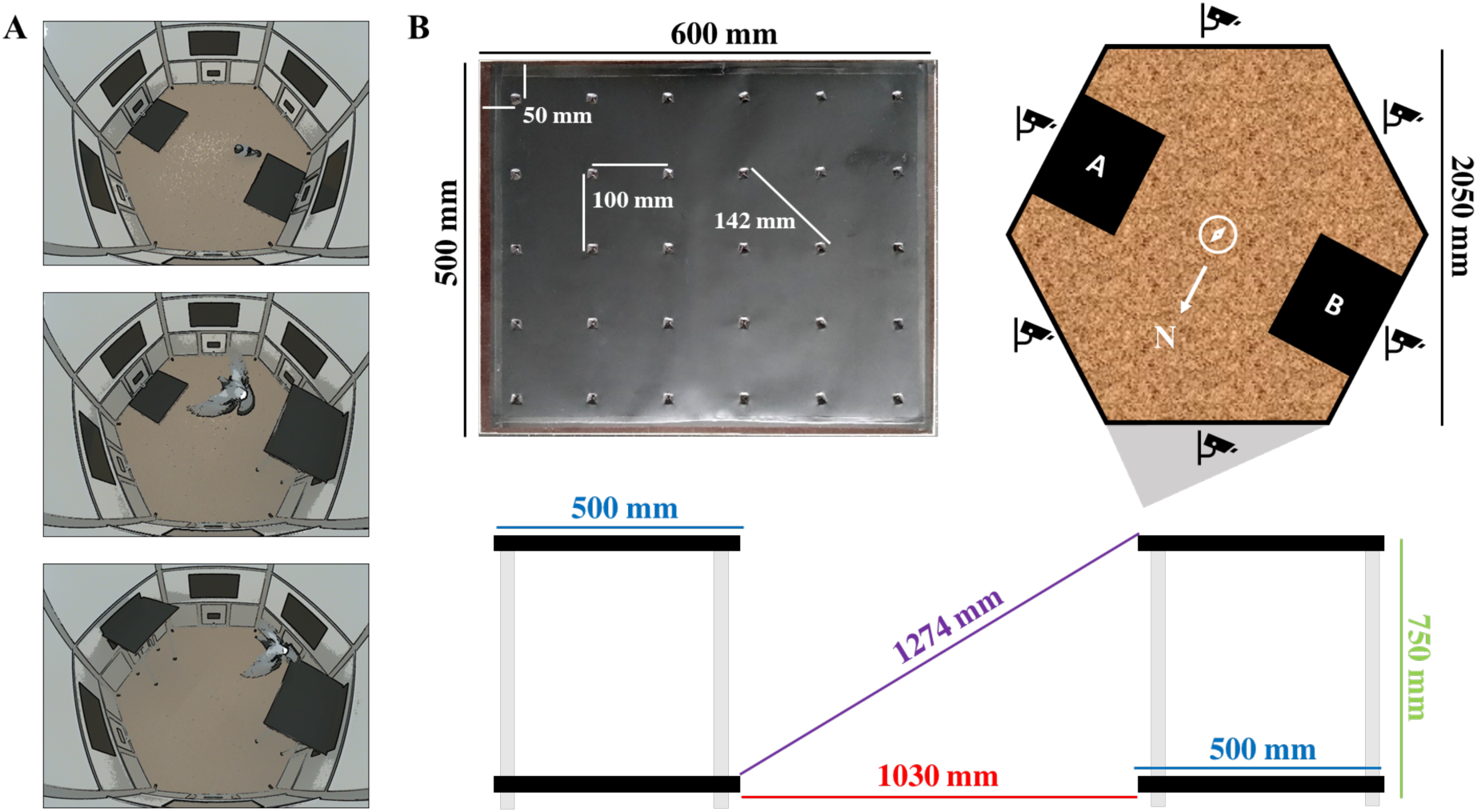
Setup of the foraging arena with two baited platforms at adjustable heights. *Note.* (A) Open field arena with two foraging platforms at different heights: 0-0 condition (top), 0-75 condition (middle) and 75-75 condition (bottom), with a pigeon transiting from platform B on the right to platform A on the left. The camera view is from above the Murphy door (see grey area on the top right of panel B) and recorded at a height of 2 meters. (B) Picture of a platform with distances between adjacent holes (top left) and schematics of the arena and platform positions (top right). Distances between platform edges (bottom). Each platform had 30 holes, 15mm wide and 15mm deep, with a single food item inside each hole.

### Procedure

During the first habituation phase, pigeons were individually placed in the arena for 20 minutes and for a minimum of four consecutive days or until they ate 85% of the food items provided in the holes of the platforms. Both platforms were placed next to each other in the center of the arena, and the food items in the platforms were visible (no plastic cover of the holes). During a second habituation phase, the platforms were covered with black adhesive film with a slit above each hole to occlude the food items and increase foraging difficulty. Pigeons were again exposed to this setup for a minimum of four 20-min sessions or until they ate 85% of the food items available.

We tested the pigeons in the three different test conditions described earlier, varying the arrangement of platform elevations, with four 20-minute sessions per condition (i.e., 12 sessions in total). Conditions were sequentially ordered for all pigeons, starting with four sessions in condition 0-0 (both platforms on the ground on opposite walls of the arena), followed by four sessions in condition 0-75 (only one of the platforms was elevated to 75 cm above the ground level), and finally four sessions in condition 75-75 (both platforms were elevated to 75 cm above the ground level). The rationale for not having counterbalanced the three conditions is that we were unsure whether the pigeons would have initially perceived an elevated platform as a place where food could be found. The position of the elevated platform was counterbalanced relative to the door in condition 0-75, and pigeons were always released in the left corner relative to the door, placing them at an equal distance from both platforms.

Pigeons were tested daily from Monday to Friday (from 9 a.m. to 4 p.m.), in the same order and at roughly the same time of the day. Data collection was split into two cohorts, where the first cohort (*n* = 6) was tested in October and November, and the second cohort was tested in February. The experimenter weighed each pigeon before and after every session to monitor the stability of individual body weights throughout the experiment and to adjust the feeding regimen, accordingly, thereby controlling for potential variability in foraging motivation between sessions.

### Data processing

Each session was video recorded from six camera angles at a 50Hz and 1440:1080px resolution, and the number of food items depleted from each platform was manually counted and recorded as session performance. We used MotionPype for camera synchronization, video processing, and behavior analysis, DeepLabCut (Mathis et al., 2018; Nath et al., 2019) for video tracking, and Anipose (Karashchuk et al., 2021) for 3D triangulation of pigeon poses. We used a Windows DELL Precision 5820 with an NVIDIA GeForce RTX 3090 40Gb GPU for video analysis, and a server computer with a 40-core Intel Xeon Gold 6248 CPU @2.50GHz and 400Gb of RAM for CPU-intensive tasks such as triangulation and data filtering. Our Python notebooks for data processing and behavior analysis, as well as the statistical analysis in R (R Core Team, 2021), are openly accessible on GitHub (see data availability statement).

Briefly, videos were compressed with HEVC/H.265 video encoding (constant rate factor crf=12) and analyzed with a pretrained DeepLabCut model for pigeons (see PigeonSuperModel in data availability statement). We applied a Viterbi filter to compute the likeliest keypoint-coordinates out of 10 predictions and smoothed the tracking replacing outliers (>20pixel offset) with a temporal median filter in a 300 ms window. The six camera views were triangulated using linear least-squares with spatio-temporal regularization implemented in Anipose to reduce reprojection, as well as temporal and spatial losses. The final 3D pose coordinates were used to extract behaviorally relevant features from time series data.

### Behavior features

The head and body positions were extracted as a spatial median centroid of a subset of key point coordinates from the pigeons’ pose, and movement kinematics as the Euclidean distance of these centroids between frames. We then classified visits to a foraging platform when the pigeons’ head was within the region of interest using the tracked platform’s edges and the head centroid. The respective timestamps were used to calculate visit lengths, latencies, and travel times between platforms. We further classified platform changes and self-transitions based on platform identity (Figure 2). We analyzed pecking behavior from time series data by localizing local minima in the head distance to the platform plane within contour lines of local oscillations and below a global median head position. We then calculated the peck count per visit, the average peck rate per second, as well as the spatial and temporal inter-peck variability as the root mean square of successive differences (RMSSD) from the time and distance between consecutive pecks. Finally, continuous behavior features per visit were aggregated and we assigned further cumulative parameters as foraging history to the current, as well as to the alternative platform on each visit, e.g. to compare the current peck rate on the current platform to the cumulative peck count throughout a session. We calculated a measure of overpecking as the ratio between the number of cumulative pecks on the current platform compared to cumulative peck count on the alternative platform, to quantify the relative value between depleting platforms.

**Figure 2.**
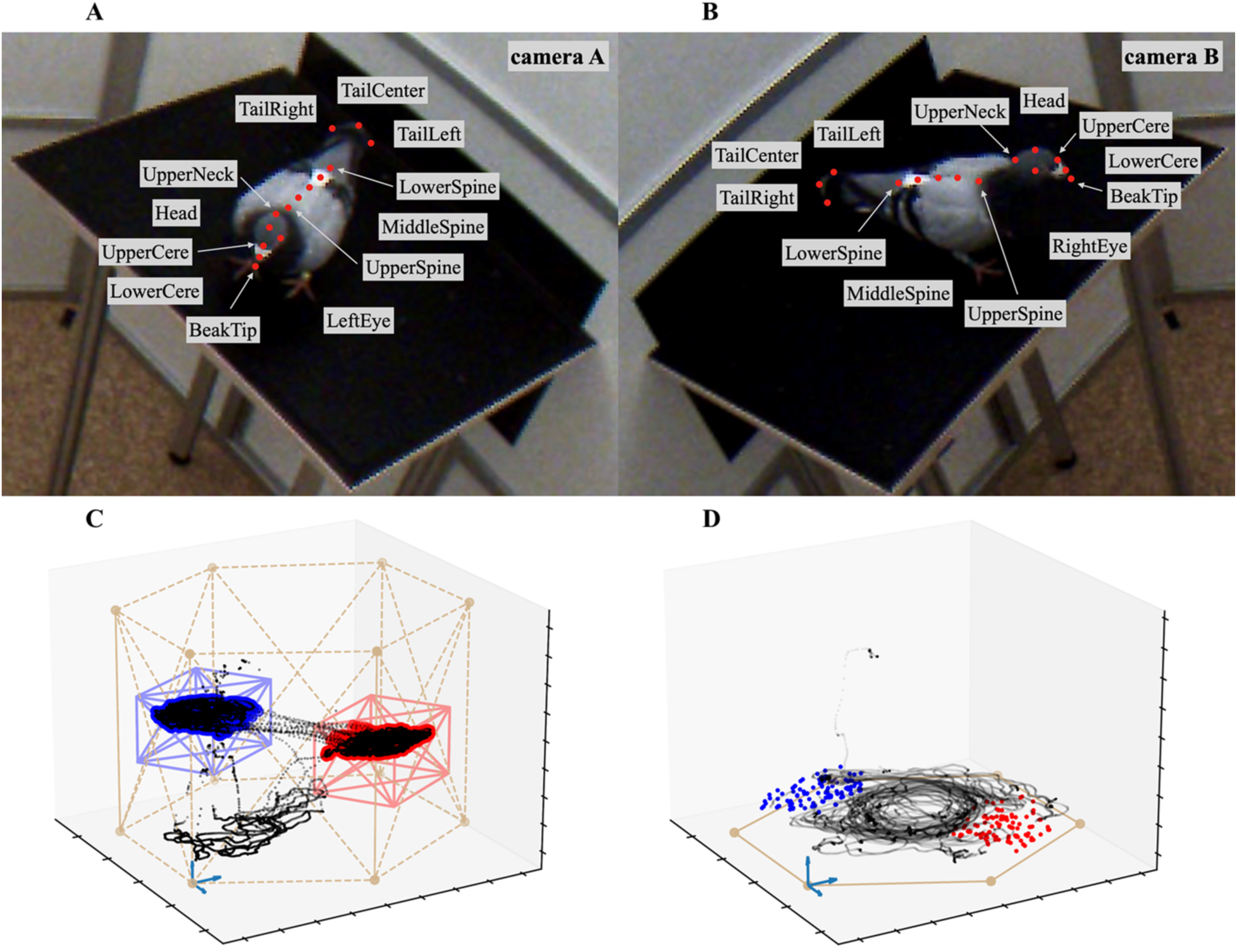
Multi-camera pose tracking and pecking detection. *Note.* (A-B) Synchronized video frames from two calibrated cameras with manually annotated keypoints. (C) 3D tracking of pigeon’s movement in the experimental arena in black, with visits to the elevated platforms marked in red and blue. (D) Pigeon’s head-movement in black, with peck-tracking on both ground platforms marked in red and blue.

### Data filtering

Three pigeons did not consume the food items during the habituation period and were excluded from the analysis (see Appendix A in the supplementary materials for individual performance curves). 44% of pigeons (4 of 9) visited the elevated platform after the first session and 67% (6 of 9) after the fourth session, in the 0-75 condition. 78% of pigeons (7 of 9) visited the elevated platform after the first session in the 75-75 condition, in which no ground alternative was available. Neither age, previous performance, nor average number of previous visits was a significant predictor for not visiting the elevated platform. Note that given the task design, failure to visit the elevated platform involved a drop in foraging performance to 50% in the 0-75 condition and to 0% in the 75-75 condition, so that platform transits could not be calculated. These individual sessions were removed from further analyses, excluding a total of 27 out of 108 test sessions due to non-visited platforms. We further trim the dataset by the highest 1% outliers in peck count per visit, visit length, and visit latency to remove possible leverage points and make later regression models more robust (see Wilcox, 2017). Data was filtered to exclude the non-foraging pigeons described above, and visits were filtered for length > 2 sec and pecks > 1.

### Statistical analysis

Unless stated otherwise, we reported median and inter-quartile ranges (IQR) instead of the mean and standard deviations to avoid leverage points in skewed and heavy tailed distributions of some of our behavior features. Given the temporal or count type of data, we expected a non-normal distribution and applied non-parametric tests such as Wilcoxon and Kruskal-Wallis, or log-transformed variables with non-normally distributed values or heteroscedastic residuals. We performed multiple linear mixed regression models with random intercept for pigeon ID to control for individual differences and repeated measures, and we included a standard set of covariates to control for age, free-feeding body weight, experimental weight, task performance, and session number. Post-hoc tests were performed for pairwise comparisons of the estimated marginal means (EMM) of each model and p-values were adjusted using the Holm method. We used the EMM as model predictions of the dependent variable for the condition parameter while adjusting for other variables in the model. This helped us calculate the pairwise comparisons for each level of the variable, as well as computing a confidence interval around the corrected estimates.

We transformed the time pigeons spent on a platform into a time-to-event problem (survival analysis) with a mixed effects Cox proportional hazard model to predict the instantaneous probability of surpassing different visit lengths. The Cox model used the proportion of visits of a certain length over time instead of the linear relationship between predictors and absolute time on platforms to estimate the instantaneous event probability of leaving a platform at any given time. To cover a wider range of not-at-risk foraging events, we augmented the dataset by 50% of censored visits. These visits were observations where the event of interest (i.e., leaving the platform) had not occurred yet, providing useful information about ongoing visits instead of focusing only on completed visits. We subset the longest 50% of visits and choose a random time point between 25-75% shorter than the actual event time to mark timepoints on visits in which pigeons did not leave the platform.

All statistical analyses were performed in R version 4.1.1, using the ‘lme4’, ‘survival’ and ‘coxme’ packages, among others (R Core Team, 2021; Bates et al. 2015; Therneau & Grambsch, 2000; Thernau, 2022).

## Results

The following results are structured in four subsections oriented on the research questions: First, we focused on pigeons’ engagement with the foraging task, analyzing latencies, side biases, and preference when choosing first to visit a platform. Second, we focused on session performance and analyzed the number of food items consumed and the number of visits per platform, comparing them between conditions as well as between platforms. Third, we took each visit as the unit of analysis for a Cox regression model to analyze the broader structure of foraging behavior, highlighting how pigeons decided to stay or leave a platform. Lastly, we closely examined each of the foraging parameters included in the previous model, to investigate how pigeons actually foraged within each visit between transitions.

### Pigeons engage with the foraging setup and postpone visits to the elevated platform

Out of our twelve pigeons, nine animals readily interacted with the experimental setup, actively foraging from the covered platforms on the ground against the walls of the arena (Figure 1). This indicated that the experiment elicited spontaneous foraging behavior with minimal habituation to the new environment. Three pigeons showed no interest in the food items used during the experiment and were excluded from the analyses (see Appendix A in the supplementary materials).

Just after placing a pigeon in the arena and closing the door, the pigeon visited the first platform with a median time latency of 10.14 seconds (IQR = 3.43), and there was no significant difference between conditions in the time to approach the first platform (Kruskal-Wallis rank sum test *X*^2^(2) = 1.90, *p* = .387). As predicted, the time latency to visit the first elevated platform was significantly shorter in the 75-75 condition (Mdn = 10.44 sec, IQR = 2.80) than in the 0-75 condition (Mdn = 122.00 sec, IQR = 109.98; Kruskal-Wallis rank sum test *X*^2^(1) = 27.43, *p* < .001). In the asymmetrical 0-75 condition, the ground platform was always visited first (two-sided binomial test, k = 15, p < .001), regardless of its horizontal position relative to the door. The elevated platform was only visited after a median delay of 105.88 seconds (IQR = 68.80). When choosing between the two platforms for the first visit, pigeons showed a slight side bias towards the platform near the door on the ground condition (0-0; two-sided binomial test, *π* = .70 (23 out of 33), *p* = .035), but the preference dissipated in the elevated condition (75-75; two-sided binomial test, *π* = .59 (16 out of 27), *p* = .442).

After visiting the first platform, pigeons moved to the second platform after a median total latency of 96.48 seconds (IQR = 75.97), with significant differences between conditions in the total time required to approach the first platform, forage from it and move to the second platform. Wilcoxon rank sum tests revealed a significantly higher total latency in condition 0-75 (Mdn = 122.00, IQR = 109.98) than in condition 0-0 (Mdn = 95.81, IQR = 71.96; *W* = 162, *p* = .044), and marginally higher total latency in condition 0-75 compared to condition 75-75 (Mdn = 89.79, IQR = 65.83; *W* = 261, *p* = .076). Pigeons showed no difference in total latency when visiting the second platforms between conditions 0-0 and 75-75 (*W* = 419, *p* = .739).

### Higher transition cost reduces number of visits but increases self-transitions

Foraging performance over an entire session, measured as the total number of food items consumed within the 20-minutes foraging time available, was consistently high across conditions, and reached ceiling levels after the first session (Figure 3). Mixed-effects analyses revealed no significant differences in session performance between conditions (all p > .05; see Model B1 in Appendix B). Comparing the foraging performance between platforms, pigeons depleted both patches evenly across conditions (Kruskal-Wallis rank sum test *X*^2^(2) = 2.97, *p* = .227), with a median difference of 0 food items (IQR = 2). In the asymmetric 0-75 condition, the pigeons depleted the platform on the ground more than the elevated platform in 6 out of 18 sessions, with a median difference of 3 food items (IQR = 6.50). Both platforms were depleted evenly in 10 out of 18 sessions, and in only 2 out of 18 sessions the elevated platform was depleted more than the ground platform by a single food item.

**Figure 3.**
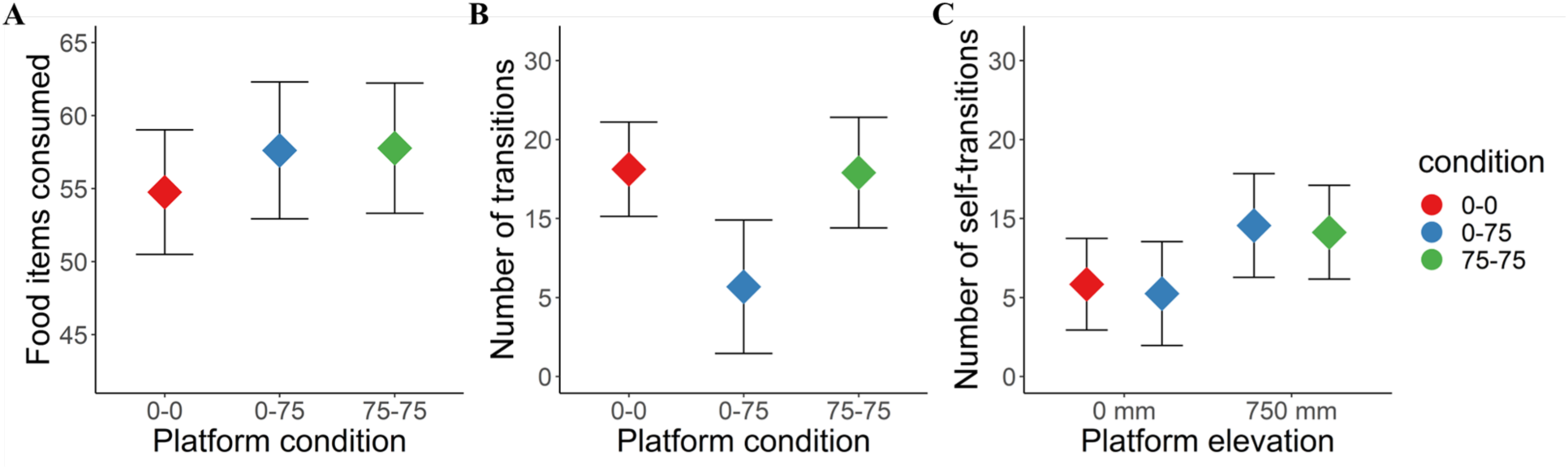
Foraging activity as platform depletion, transitions, self-transitions between conditions. *Note.* Estimated marginal means (adjusted for covariates in mixed regression models) with 95% confidence intervals for foraging performance of 9 pigeons across 12 consecutive sessions under three platform configurations: both platforms at ground level (0–0), one platform elevated to 750 mm (0–75), and both platforms elevated (75–75). (A) Number of food items consumed per session. (B) Number of transitions between platforms. (C) Number of self-transitions (departures and returns to the same platform).

To forage a total of 60 food items from the two platforms, pigeons performed a median of 12 transitions (IQR = 15) per session. A linear mixed effects regression model confirmed that the number of transitions between platforms within a session differed significantly between condition, with significantly less transitions in condition 0-75 compared to conditions 0-0 (*b* = 10.26, *t* = 3.00, *p* = .004) and 75-75 (*b* = 9.96, *t* = 2.79, *p* = .014). There was no difference in the number of transitions between ground platforms in condition 0-0 and elevated platforms in condition 75-75 (*b* = 0.30, *t* = 0.10, *p* = .921), and the number of transitions between platforms within a session was significantly reduced by the pigeons’ age (*b* = −4.20, *t* = 3.14, *p* = .044; Model B2).

Apart from changing platforms, pigeons also performed a median of 10 self-transitions (IQR = 10) per session. In other words, they left the current platform, usually to recover a lost food item, and returned to the same platform instead of moving to the other one. A linear mixed effects regression model showed that the number of self-transitions was significantly higher on the elevated platform of the 0-75 condition than on the ground platform of the same condition (*b* = 5.94, *t* = 2.96, *p* = .011), and higher compared to the ground platforms of the 0-0 condition (*b* = 5.13, *t* = 3.03, *p* = .011). The number of self-transitions to the elevated platforms of the 75-75 condition was also significantly higher than to the ground platforms of the 0-0 condition (*b* = 4.53, *t* = 3.79, *p* = .001) but did not differ from the number of self-transitions to the other elevated platform of the 0-75 condition (*b* = 0.59, *t* = 0.36, *p* = .999). Similarly, the ground platform of the 0-75 condition had significantly less self-transitions than the elevated platforms of the 75-75 condition (*b* = −5.35, *t* = 3.21, *p* = .008) but did not differ from the ground platforms in the 0-0 condition (*b* = −0.82, *t* = 0.48, *p* = .999). No other parameter in the model had a significant effect (Model B3, Figure 3).

Throughout a foraging session, pigeons spent cumulatively a median of 527.52 seconds (IQR = 340.78) or 43.96% of the session on platforms and 89.62 seconds (IQR = 108.42) or 7.47% traveling. Comparing these figures over time, during the first 10 minutes they spent 50.93% of time on platforms, and only 25.02% during the last 10 minutes of a session.

### Modeling the structure of behavior: Stay-or-leave decisions are predicted by foraging history and transition cost

We analyzed visit duration as time-to-event data (i.e. latency to leave) using a Cox proportional hazards model to estimate the instantaneous probability of leaving a platform. Predictors included platform condition, current foraging parameters, and cumulative foraging history (Table 1, Figure 4). Predictors were entered hierarchically to test the effect of each group of parameters independently, and model fit improved with each step. Adding current foraging parameters (Step 2) and cumulative foraging history (Step 3), each contributed significantly to the prediction of foraging decisions. While the effects of individual parameters are discussed further, it is sufficient to note here that incorporating these variables in a multivariate model substantially enhanced predictive accuracy. Full model parameters and regression coefficients are presented in Table 1.

**Figure 4.**
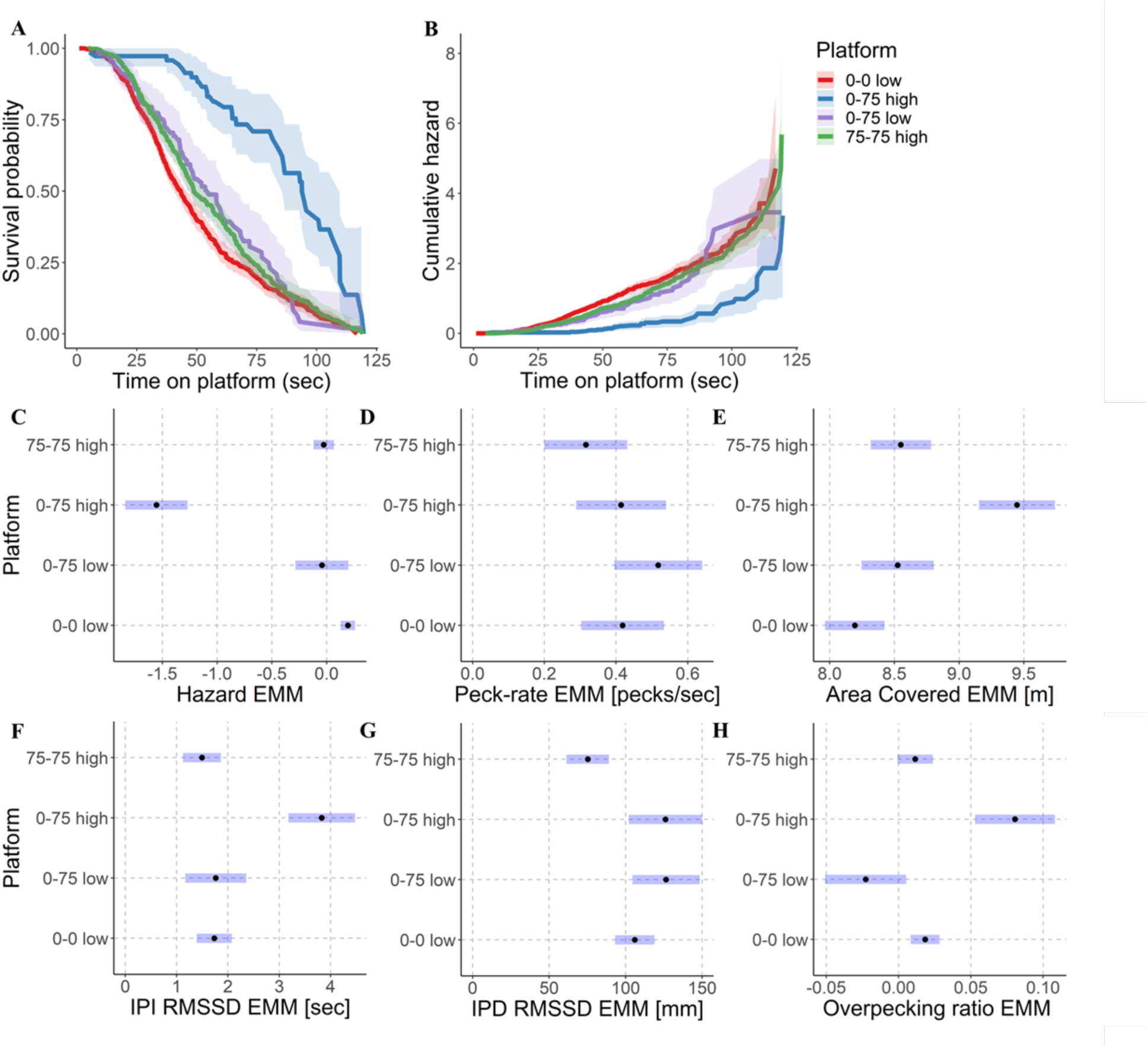
Cox Regression Model on the instantaneous probability of leaving a platform. *Note.* Results from Cox regression and follow-up models examining platform occupancy behavior. (A) Survival probability, indicating the likelihood of a pigeon remaining on a platform after a given time. (B) Cumulative hazard, indicating the likelihood of having left a platform after a given time. (C) Adjusted mean hazard values for each condition from the Cox regression model. (D–H) Adjusted mean values for each condition from separate regression models for peck rate (D), total area covered on the platform (E), inter-peck interval variability (IPI RMSSD; F), inter-peck distance variability (IPD RMSSD; G), and overpecking ratio (H). Error bars or shaded regions represent 95% confidence intervals.

**Table 1.**
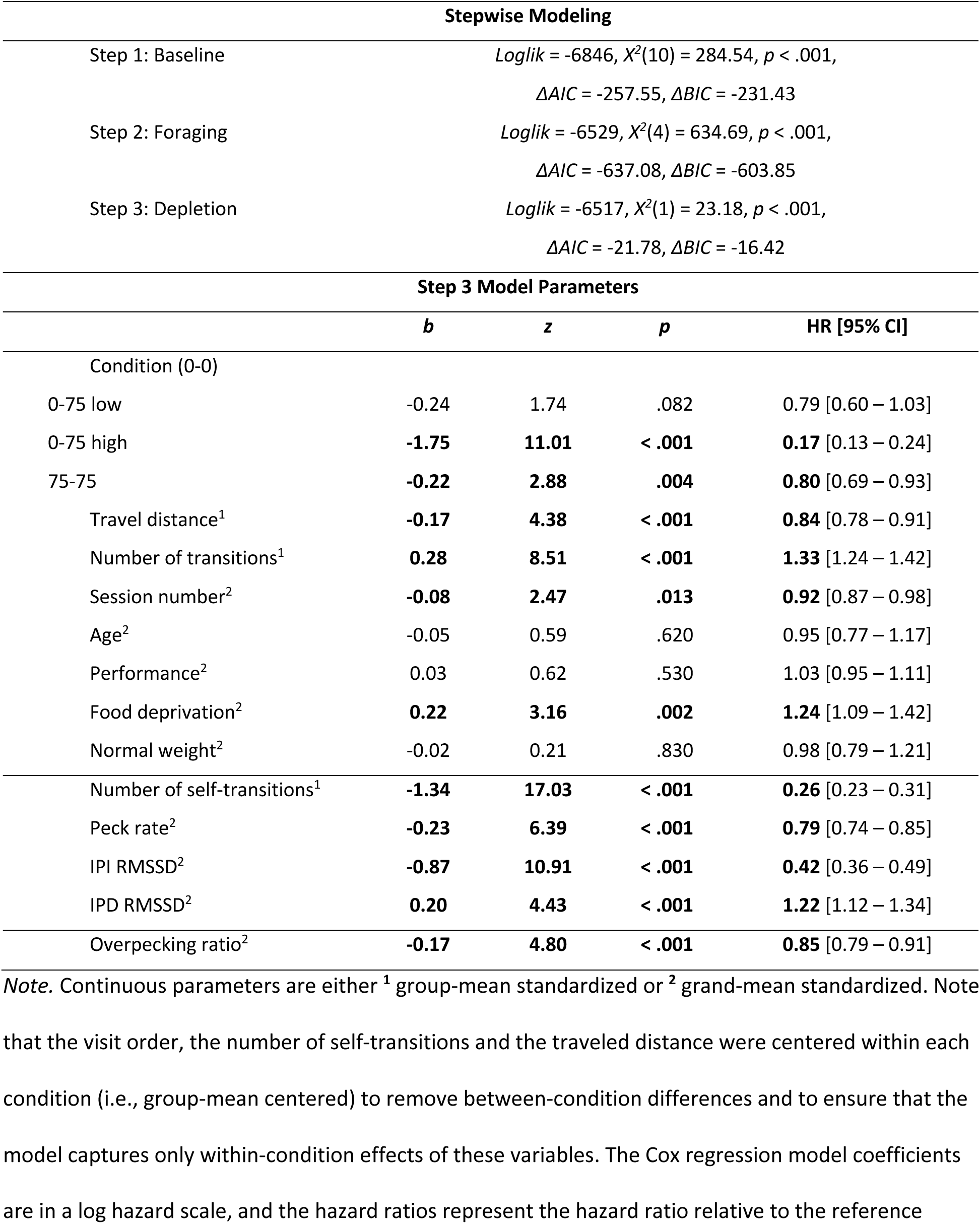
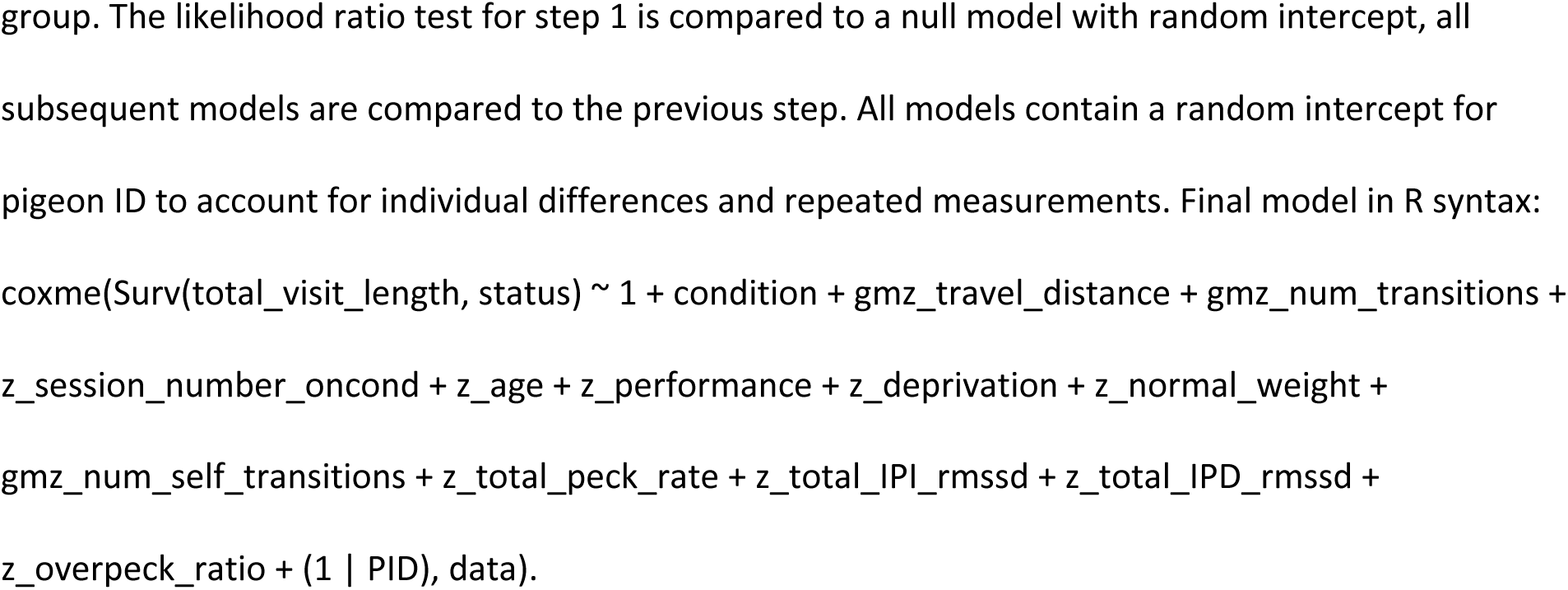
Stepwise Cox Regression Models for probability to leave platforms during foraging visits.

| Stepwise Modeling |  |  |  |  |
| --- | --- | --- | --- | --- |
| Step 1: Baseline | $Loglik = -6846, X^2(10) = 284.54, p < .001,$<br>$\Delta AIC = -257.55, \Delta BIC = -231.43$ | | | |
| Step 2: Foraging | $Loglik = -6529, X^2(4) = 634.69, p < .001,$<br>$\Delta AIC = -637.08, \Delta BIC = -603.85$ | | | |
| Step 3: Depletion | $Loglik = -6517, X^2(1) = 23.18, p < .001,$<br>$\Delta AIC = -21.78, \Delta BIC = -16.42$ | | | |
| Step 3 Model Parameters |  |  |  |  |
|  | *b* | *z* | *p* | HR [95% CI] |
| Condition (0-0) |  |  |  |  |
| 0-75 low | -0.24 | 1.74 | .082 | 0.79 [0.60 – 1.03] |
| 0-75 high | **-1.75** | **11.01** | **< .001** | **0.17** [0.13 – 0.24] |
| 75-75 | **-0.22** | **2.88** | **.004** | **0.80** [0.69 – 0.93] |
| Travel distance<sup>1</sup> | **-0.17** | **4.38** | **< .001** | **0.84** [0.78 – 0.91] |
| Number of transitions<sup>1</sup> | **0.28** | **8.51** | **< .001** | **1.33** [1.24 – 1.42] |
| Session number<sup>2</sup> | **-0.08** | **2.47** | **.013** | **0.92** [0.87 – 0.98] |
| Age<sup>2</sup> | -0.05 | 0.59 | .620 | 0.95 [0.77 – 1.17] |
| Performance<sup>2</sup> | 0.03 | 0.62 | .530 | 1.03 [0.95 – 1.11] |
| Food deprivation<sup>2</sup> | **0.22** | **3.16** | **.002** | **1.24** [1.09 – 1.42] |
| Normal weight<sup>2</sup> | -0.02 | 0.21 | .830 | 0.98 [0.79 – 1.21] |
| Number of self-transitions<sup>1</sup> | **-1.34** | **17.03** | **< .001** | **0.26** [0.23 – 0.31] |
| Peck rate<sup>2</sup> | **-0.23** | **6.39** | **< .001** | **0.79** [0.74 – 0.85] |
| IPI RMSSD<sup>2</sup> | **-0.87** | **10.91** | **< .001** | **0.42** [0.36 – 0.49] |
| IPD RMSSD<sup>2</sup> | **0.20** | **4.43** | **< .001** | **1.22** [1.12 – 1.34] |
| Overpecking ratio<sup>2</sup> | **-0.17** | **4.80** | **< .001** | **0.85** [0.79 – 0.91] |

*Note.* Continuous parameters are either <sup>1</sup> group-mean standardized or <sup>2</sup> grand-mean standardized. Note
that the visit order, the number of self-transitions and the traveled distance were centered within each condition (i.e., group-mean centered) to remove between-condition differences and to ensure that the model captures only within-condition effects of these variables. The Cox regression model coefficients are in a log hazard scale, and the hazard ratios represent the hazard ratio relative to the reference
group. The likelihood ratio test for step 1 is compared to a null model with random intercept, all subsequent models are compared to the previous step. All models contain a random intercept for pigeon ID to account for individual differences and repeated measurements. Final model in R syntax:
``` coxme(Surv(total_visit_length, status) ~ 1 + condition + gmz_travel_distance + gmz_num_transitions + z_session_number_oncond + z_age + z_performance + z_deprivation + z_normal_weight + gmz_num_self_transitions + z_total_peck_rate + z_total_IPI_rmssd + z_total_IPD_rmssd + z_overpeck_ratio + (1 | PID), data). ```

As hypothesized, the final model showed a significant main effect of platform condition or travel modality on the timing of visits and transitions (*X^2^*(3) = 121.91, *p* < .001). Specifically, pairwise post-hoc comparisons with Holm correction revealed a significantly lower probability of leaving the elevated platform in the 0-75 condition than in the 75-75 condition (*b* = 1.53, *z* = 9.45, *p* < .001), and a lower probability of leaving a platform in the elevated 75-75 condition than in the ground 0-0 condition (*b* = 0.22, *z* = 2.88, *p* = .012). Within the 0-75 condition, the probability to leave the ground platform was significantly higher than the probability to leave the elevated platform (*b* = 1.51, *z* = 8.02, *p* < .001). The probability of leaving the elevated platform in the 0-75 condition was also significantly lower than in the 0-0 condition (*b* = 1.75, *z* = 11.01, *p* < .001). There were no significant differences in the probability of leaving a platform between the ground platform of the 0-75 condition and 0-0 condition (*b* = −0.24, *z* = 1.74, *p* = .164), or between the ground platform of the 0-75 and 75-75 condition (*b* = 0.02, *z* = 0.11, *p* = .911).

The Cox regression model in Table 1 also showed a significant effect of the increasing number of previous transitions, i.e. the re-visiting order, on the probability of leaving a platform (HR = 1.33 95%-CI [1.24 – 1.42], *b* = 0.28, *z* = 8.51, *p* < .001), effectively shortening the visit length. The session number for each condition, i.e. the number of repeated exposures to a given platform arrangement, significantly reduced the probability of leaving a platform (HR = 0.92 95%-CI [0.87 – 0.98], *b* = −0.08, *z* = 2.74, *p* = .013). Similarly, a higher number of self-transitions within a visit reduced the probability of leaving a platform (HR = 0.26 95%-CI [0.23 – 0.31], *b* = −1.34, *z* = 17.03, *p* < .001). On the other hand, pigeons with a lower experimental weight relative to their free-feeding body weight had a higher probability to leave the platform at any given time (HR = 1.24 95%-CI [1.08 – 1.42], *b* = 0.22, *z* = 3.16, *p* = .002). Furthermore, the traveled distance towards a platform had a significant effect on the probability to leave a platform, HR = 0.84 95%-CI [0.78 – 0.91], *b* = −0.17, *z* = 4.38, *p* < .001, with longer traveled distances towards a platform reducing the probability to leave, and in turn extending the current visit length. Full model coefficients are provided in Table 1, and Figure 4 illustrates survival probabilities and pairwise comparisons between platform conditions.

### Reduced foraging activity over time, and higher variability with increasing transition cost

We next examined individual foraging parameters to characterize within-visit behavior across conditions. In the analysis above, we assumed platform visits and a certain time-on-platform to be directly related to foraging activity, but we further analyzed the pigeons’ head movement with 3D tracking to measure foraging behavior more directly. Specifically, we analyzed both the horizontal head movement foraging on the platform, i.e. search area, as well as the number of vertical pecks towards the platform, the temporal inter-peck-intervals (IPI), the spatial inter-peck-distances (IPD), and the overall root mean square of successive differences (RMSSD) variability of pigeons’ pecking behavior (Figure 2).

On average, we registered a median of 222.5 pecks per session (IQR = 230.5) in the 0-0 condition, a median of 238.5 pecks (IQR = 282.75) in the 0-75 condition, and a median of 137 pecks (IQR = 158) in the 75-75 condition. The observed number of pecks was much larger than the 60 pecks normally required (60 holes were available). As a cumulative parameter, the total number of pecks per visit was strongly correlated with the visit length (Pearson’s *r* = 0.67, *t* = 29.59, *p* < .001), and we therefore used the peck rate per second as a standard measure, instead of the absolute peck count. Nevertheless, the absolute peck count was significantly negatively correlated with visit order (Pearson’s *r* = −0.32, *t* = 11.38, *p* < .001).

The Cox regression model in Table 1 showed a significant effect of the current peck rate, reducing the probability to leave a platform, HR = 0.79 95%-CI [0.74 – 0.85]. With the peck rate as a measure of foraging intensity, we had predicted an increasing pecking rate with decreasing food density, i.e. with an increasing number of previous visits in a session. Instead, a linear mixed regression model showed a significant decrease in peck rate with an increasing number of visits (*b* = −0.07, *t* = 9.10, *p* < .001), as well as a higher overall peck rate for pigeons with higher levels of food deprivation (*b* = 0.05, *t* = 2.63, *p* = .009). Between conditions, the 75-75 platforms had a significantly lower peck rate than the ground platforms in the 0-0 condition (*b* = 0.10, *t* = 5.42, *p* < .001), and the ground platform in the 0-75 condition (*b* = 0.20, *t* = 5.81, *p* < .001), as well as the elevated platform in the 0-75 condition (*b* = 0.10, *t* = 2.59, *p* = .030). The ground platform in the 0-75 condition had a higher peck-rate than the ground platforms in the 0-0 condition (*b* = 0.10, *t* = 2.87, *p* = .017), and a marginally higher peck-rate than on the elevated platform in the 0-75 condition (*b* = 0.10, *t* = 2.22, *p* = .054). The peck rate associated with the elevated platform in the 0-75 condition and the ground platforms in the 0-0 condition did not differ (*b* = 0.004, *t* = 0.11, *p* = .915; Figure 4d, Model B4).

We further predicted an increase in pecking variability over time, since the few remaining food items might be more sparsely distributed after the initial visits. The Cox Regression model in Table 1 showed significant effects of pecking variability on the probability to leave a platform. Pigeons showed a lower probability of leaving the platform when foraging with a higher IPI RMSSD, HR = 0.42 95%-CI [0.36 – 0.49], and a lower IPD RMSSD, HR = 1.22 95%-CI [1.12 – 1.34]. In separate mixed regression models for each of the parameters, the RMSSD variability of IPI decreased with an increasing number of previous visits (*b* = −0.20, *t* = 3.00, *p* = .003), as well as with a higher session performance (*b* = −0.23, *t* = 2.76, *p* = .007). It was significantly higher on the elevated platform in the 0-75 condition compared to conditions 0-0 (*b* = 2.09, *t* = 6.42, *p* < .001), condition 75-75 (*b* = 2.33, *t* = 7.10, *p* < .001), and compared to the ground platform in condition 0-75 (*b* = 2.06, *t* = 5.11, *p* < .001). The other comparisons in the model were not significant (Figure 4f, Model B5). A second linear mixed regression showed a decrease in RMSSD of IPD with ascending visit order within a session (*b* = −21.96, *t* = 9.20, *p* < .001), as well as a marginal decrease with increasing age of the pigeon (*b* = −11.77, *t* = 2.33, *p* = .055). Pigeons foraged from the elevated platforms in condition 75-75 with a significantly lower IPD RMSSD than on the ground platforms of both the 0-0 condition (*b* = −30.74, *t* = 5.21, *p* < .001) and 0-75 condition (*b* = −51.18, *t* = 4.65, *p* < .001). Further, the IPD RMSSD on the elevated platforms in the 75-75 condition was significantly lower than on the elevated platform in the 0-75 condition (*b* = −50.82, *t* = 4.17, *p* < .001; Figure 4g, Model B6).

We also predicted higher area coverage in conditions with higher transition cost between platforms, and a linear mixed regression model on the log-transformed head distance on a platform confirmed that the area covered searching on each platform significantly decreased with increasing number of previous visits (*b* = −0.32, *t* = 13.53, *p* < .001). This result indicates decreasing foraging activity proportional to the decreasing rate of food intake due to patch depletion. Pigeons covered a significantly larger area per visit on the ground platform in the 0-75 condition than in the 0-0 condition (*b* = 0.33, *t* = 3.01, *p* = .005), and a larger area on the elevated platform in the 0-75 condition compared to the 0-0 condition (*b* = 1.25, *t* = 10.43, *p* < .001), as well as the ground platform in the 0-75 condition (*b* = 0.92, *t* = 6.22, *p* < .001). The distance covered searching on the elevated platform in condition 0-75 was also significantly larger than in the 75-75 condition (*b* = 0.90, *t* = 7.44, *p* < .001). Furthermore, pigeons covered a significantly larger area per visit on the elevated platforms in condition 75-75 than in the 0-0 condition (*b* = 0.36, *t* = 5.95, *p* < .001), but not more than on the ground platform in the 0-75 condition (*b* = 0.02, *t* = 0.22, *p* = .824; Figure 4e, Model B7). These results reflect increased investment on the current platform relative to the increased transition cost between and towards elevated platforms.

In the MVT, the habitat average describes the expected rate of food intake across all remaining patches in the environment, but considering only two food patches, we quantified relative cumulative foraging between platforms as an overpecking ratio. We defined overpecking as the ratio of cumulative pecks on the current platform relative to the alternative platform within a session. An overpecking ratio near zero indicates relatively equal cumulative foraging on both platforms, either both still pristine with a high food density or both equally depleted towards the end of a foraging session. A positive overpecking ratio indicates that the current platform had been depleted more than the alternative, and a negative overpecking ratio indicates that the current platform had been depleted less than the alternative and was therefore left under-foraged.

The foraging behavior model in Table 1 showed a significant effect of the overpecking ratio on the probability of leaving a platform, with a higher overpecking ratio reducing the probability of leaving the current platform, HR = 0.85, 95%-CI [0.79 – 0.91]. It is important to note that overpecking refers to the ratio between platforms at the end of a visit, and higher overpecking was overall moderately correlated with longer visits (Pearson’s *r* = .36, *t* = 12.81, *p* < .001). We investigated this further with a separate linear mixed effects model and found that the overpecking ratio significantly decreased with an increasing number of previous visits to a platform (*b* = −0.01, *t* = 2.13, *p* = .033), as well as with higher age of the pigeon (*b* = −0.01, *t* = 2.40, *p* = .017). More interestingly, the overpecking behavior differed significantly between conditions. Pigeons overpecked the elevated platform in condition 0-75 significantly more than the ground platform in the same condition 0-75 (*b* = 0.10, *t* = 5.27, *p* < .001).

Similarly, the elevated platform of condition 0-75 was overpecked more than the platforms in conditions 0-0 (*b* = 0.06, *t* = 4.20, *p* < .001) and 75-75 (*b* = 0.07, *t* = 4.57, *p* < .001). Platforms in conditions 0-0 and 75-75 were overpecked similarly (*b* = 0.01, *t* = 0.93, *p* = .352), and the ground platform in condition 0-75 was overpecked less than platforms in conditions 0-0 (*b* = −0.04, *t* = 2.73, *p* = .020) and 75-75 (*b* = −0.03, *t* = 2.23, *p* = .051). Overall, these differences show the highest over-foraging tendency on the elevated platform of the 0-75 condition, followed by the 75-75 and the 0-0 condition, and finally the ground platform of the 0-75 condition (Figure 4h, Model B8). The session number, performance, level of food deprivation or normal body weight were not significantly related to overpecking.

## Discussion

In this study, we examined the foraging behavior of pigeons in a controlled laboratory environment designed to approximate semi-natural conditions. Using a custom-built apparatus with spatially distinct foraging platforms, we manipulated the cost of travel between patches by varying their elevation, thereby comparing bipedal locomotion with short flights. Our results show that a 75 cm elevation difference between platforms (0-75 condition), compared to symmetric configurations (0-0 and 75-75 conditions), significantly altered foraging patterns in pigeons, increasing the cost of travel between patches and affecting platform choice, visit duration, and overall foraging activity in ways consistent with predictions from the MVT (Charnov, 1976). Specifically, the differences in foraging activity across conditions and platforms suggest that (a) transition costs are a meaningful determinant of patch choice, and (b) these costs can be estimated and inferred from consistent overpecking and patch depletion patterns. In this respect, our findings are also consistent with those reported by Kono (2019), showing that pigeons forage optimally in situations of diminishing returns (see also Hanson & Green, 1989; Wanchisen et al., 1988).

Our experimental setup (Figure 1) successfully elicited spontaneous, naturalistic foraging behavior in pigeons while maintaining experimental control. Using automated 3D video tracking provided high-resolution data on body movements and pecking behavior, allowing for non-invasive, detailed quantification of spatial and temporal foraging patterns (Figure 2). This computational approach not only enhanced measurement precision but also offers a modular framework that can be adapted for tracking other behaviors, such as head direction in social contexts (see Delacoux & Kano, 2024). Finally, our use of a multivariate Cox proportional hazards model revealed that pigeons’ patch-leaving decisions reflect relative changes in marginal value of food patches, supporting the idea that foraging strategies emerge dynamically from the interaction between internal states and environmental structure. In the following sections, we interpret these findings in light of our initial predictions and the broader context of foraging theory.

### Pigeons favor the platform associated with minimal travel cost

We found that pigeons consistently visited the ground platform first in the asymmetric elevation condition (0-75), regardless of its lateral position. This pattern suggests a clear preference for minimizing travel effort when options differ. In the symmetric 0-0 condition, pigeons showed a slight side bias toward the platform closest to the door, but this bias disappeared in the 75-75 condition, where both options were elevated. Notably, the latency to approach the first platform did not differ between the 0-0 and 75-75 conditions, despite the added elevation cost in the latter, suggesting that pigeons respond similarly when both options impose equivalent energetic demands. These findings align with the predictions of Charnov’s (1976) MVT: When the transit costs between patches differ, animals should favor the lower-cost option to maximize net gain.

In line with this, we also found that the distance traveled toward a platform significantly affected the probability of leaving that platform. Specifically, longer travel distances were associated with longer visit durations, consistent with the idea that greater investment increases the threshold for departure. Importantly, this effect of travel distance was statistically independent from elevation and remained significant after controlling for task parameters using a Cox proportional hazards model. Together, these results show that pigeons take both distance and locomotion costs into account when making foraging decisions, further validating MVT under semi-naturalistic, dynamic conditions.

### Travel cost increases transition latency and patch residence time

On the first visit of a session, pigeons spent more time on the ground platform in the 0-75 condition compared to either platform in the 0-0 and 75-75 conditions. The latency before starting the second visit was significantly higher in the 0-75 condition than in the symmetric conditions, indicating that the added vertical distance imposed a measurable travel cost. In contrast, we found no difference in transition latency between the 0-0 and 75-75 conditions, despite the need for short flights in the latter. This absence of a difference may be due to increasing familiarity with the setup across sessions and the influence of condition order, as pigeons shortened their transition times with experience. In stable environments, like the one we used, a variety of species are known to adjust foraging strategies based on past experience and perceived patch quality (e.g., Berger-Tal et al., 2014; Kamil and Roitblat, 1985; Nakata et al., 2003; Sabrina et al., 2014).

The longer residence time on the ground platform in the 0-75 condition during first visit may reflect a behavioral adjustment to increased patch-leaving costs. According to the MVT, an animal should leave a patch when the rate of food intake falls below the average rate expected from the alternative (Charnov, 1976; Stephens & Krebs, 1986; Todd & Kacelnik, 1993). When travel costs are higher, the expected net gain from switching patches is reduced, thereby lowering the marginal value threshold and increasing the time spent foraging at the current site. In our setup, although the second patch offered the same food quantity across conditions, access to the elevated platform in the 0-75 condition required greater effort, explaining the extended patch residence times observed on the ground patch.

Using a Cox proportional hazards model controlling for session number and visit order, we analyzed all platform visits and revisitations throughout foraging sessions. Interestingly, this analysis revealed that pigeons spent more time on the elevated platform in the 0-75 condition than on any other platforms, a reversal from the initial visit pattern above, where the initial visit to the ground platform had longer residence times. Similarly, visits to elevated platforms in the 75-75 condition were longer than those to ground platforms in the 0-0 condition. This shift may reflect a dynamic adjustment in patch use over time: while pigeons initially favor lower-cost patches to minimize early effort, repeated visits and patch depletion likely increase the relative value of elevated platforms, especially once the higher travel costs have been invested. These findings may highlight that patch residence time depends not only on immediate travel costs but also on experience, patch depletion, and possibly perceived safety or comfort associated with elevation, factors that all align with MVT’s predictions under dynamic conditions.

### Elevated platforms reduce patch switching and increase self-transitions

Although total food consumption did not differ across conditions, pigeons made fewer transitions between platforms in the 0-75 condition compared to both 0-0 and 75-75 conditions, which were equivalent in this respect. However, elevated platforms in both the 0-75 and 75-75 conditions showed significantly more self-transitions, which denote brief departures and returns to the same platform without any visit to the alternative. We believe this behavioral pattern does not necessarily reflect stronger motivation to stay on elevated patches with pigeons tested individually rather than collectively (e.g., Portugal et al., 2017). Instead, elevated platforms had discrete edges and limited maneuvering space, making it more difficult for pigeons to reach food items near the periphery without stepping off. In contrast, ground platforms were better integrated into the environment: pigeons could stretch their necks to reach items beyond the platform edge or walk off while continuing to peck without leaving the platform.

### Foraging activity declines with patch depletion

The number of previous visits to a platform significantly increased the probability of leaving it sooner, reflecting diminishing returns with repeated visits. Similarly, the area covered on each platform, as well as the RMSSD variability of IPI and IPD, decreased as visit number increased. Pecking intensity (peck rate) and the absolute number of pecks per visit also declined steadily throughout the session.

The predicted increase in pecking distribution with decreasing food density was not confirmed via spatio-temporal variability in inter-peck intervals and inter-peck distances. Instead, pigeons showed a tendency toward more stable and homogeneous foraging patterns. These findings seem to contradict previous evidence that pigeons increased the number of pecks and visit duration in poorer compared to richer areas, specifically in contexts where baited holes were inconsistently versus consistently rewarded (Anselme et al., 2022, 2024; see also Hanson & Green, 1989). However, several factors may account for these differences. In the present study, the number of holes per platform was smaller, session duration was longer, and all holes were baited equally at the start. As such, there was no initial uncertainty and little temporal pressure for foraging, which may have reduced the need for extensive searching. This likely contributed to the overall reduction in area covered during later visits. In contrast, studies using larger platforms with variable baiting have shown increased inter-peck distance variability, likely reflecting uncertainty and missed food (Wittek et al., 2022).

Later in the session, pigeons spent more time off-platform, switching from active foraging to exploratory behavior, coinciding with patch depletion. This behavior resembles what pigeons would do in the wild after exploiting a known food source, an option not available in the abovementioned studies in which no space was available around the platform. In our setup, the opportunity to leave the foraging patches may have allowed a more natural shift toward exploration, rather than continued overexploitation.

### Ongoing foraging behavior predicts patch-leaving decisions

Our findings demonstrate that pigeons’ foraging decisions can be meaningfully predicted from their own ongoing behavior, rather than from objective reinforcement rates—consistent with evidence accumulation models of decision-making (e.g., Davidson & Hady, 2019; Piet et al., 2018; Zhang & Hui, 2014). By modeling behavioral indicators such as peck rate, IPI, and IPD, we captured how pigeons adjust their patch-leaving behavior in response to local intake conditions (see Table 1). Importantly, our model showed that decisions are influenced not only by the immediate state of a patch but also by its relative cost and value within the broader foraging environment. The model’s predictions improved with the inclusion of the overpecking ratio, emphasizing how pigeons’ behavior reflects sensitivity to relative returns, without requiring an explicit internal representation of cumulative value.

Consistent with this, higher overpecking was associated with a lower probability of leaving, suggesting that cumulative foraging investment serves as a proxy for perceived patch value. Interestingly, overpecking was the highest on the elevated platform of the 0-75 condition and the lowest (i.e., under-pecking) on its ground counterpart, despite expectations of greater foraging activity on the ground platform before transitions. No significant differences in overpecking emerged between the symmetric conditions (0-0 and 75-75). This asymmetry may reflect increased effort or physical constraints on elevated platforms, which often required more self-transitions, broader spatial coverage, and greater temporal variability (IPI RMSSD).

Alternatively, pigeons may overexploit the elevated patch once the travel cost is sunk, reducing the likelihood of revisiting it later, consistent with the lower number of total transitions observed in this condition. Finally, the overpecking ratio declined with repeated visits, suggesting that pigeons adjusted their departure threshold over time, either because of patch depletion or reduced reinforcement relative to previous expectations.

### Individual differences in motivation and foraging behavior

Although age and body weight were not experimentally manipulated, our sample showed some differences in foraging behavior associated with these factors. Older pigeons made fewer transitions between platforms and showed slightly lower variability in inter-peck distances and reduced overpecking, possibly reflecting more stable or efficient (experience-based) foraging behavior (e.g., Weimerskirch et al., 2005). Pigeons with a lower experimental weight relative to their free-feeding body weight tended to leave platforms earlier and pecked at a higher rate, suggesting an elevated motivational state. These results should be interpreted with caution, as age was not normally distributed and the targeted experimental body weights were relatively homogeneous across individuals. Nevertheless, such individual differences may reflect natural variations in motivation and cognition with significant effects on foraging, whether in the lab or in the wild (Flaim & Blaisdell, 2023; Wittek et al., 2024).

### Limitations and potential biases

This study has some limitations that may affect the generalizability of our findings. The sample size was relatively small, composed of a single gender, and featured limited variability in age, body weight, and food deprivation levels (after excluding 3 non-engaging animals). Additionally, the simplicity of the task, with only two closely positioned food platforms, may have limited the opportunity to observe more nuanced differences in foraging behavior. The cost of travel was moderate but likely not high enough to elicit robust differences between bipedal locomotion and short flight, and the physical scale of the foraging environment may not fully capture the complexity of more naturalistic foraging scenarios.

Several factors in the design may have introduced unintended biases. Elevated platforms may have served both as a cost and as a safe vantage point, potentially affecting patch preference independently of travel effort. Self-transitions, sometimes due to dropped food or limited maneuverability on elevated platforms, were aggregated into single visits, possibly conflating different behavioral patterns. Additionally, the salience of only two visible options eliminated any cost associated with environmental search, reducing the ecological realism of the decision-making context. Finally, assumptions such as rapid familiarity with platforms and patch quality, as well as reduced perceived risk in a controlled arena, may not fully reflect natural foraging dynamics. Future studies could improve ecological validity by introducing more patches with variable visibility, including decoy platforms, disambiguating elevation from perceived safety, and continuously tracking behavior to better estimate marginal value over time. The proposed setup also lends itself well to comparative studies across species, including those with different ecological foraging strategies such as caching.

### Conclusion

This study demonstrates the value of combining advanced tracking and modeling techniques to investigate animal behavior in a naturalistic but controlled setting. We used advanced 3D video tracking to track head and body movements and to quantify foraging activity at a fine scale without invasive equipment. Combined with a Cox regression approach to model decisions based on observed behavior over time, this method offers a practical framework for studying decision-making in freely moving animals. It can be extended to other species, tasks, or setups where the timing of decisions and the structure of the environment are central to behavior.

Building on these methodological advances, our findings show that pigeons adapt their foraging behavior based on the physical effort required to move between food patches and on their own ongoing activity within each patch. By introducing differences in travel conditions through platform elevation and distance, we were able to alter pigeons’ choices, visit durations, and movement patterns in ways consistent with the MVT. These results suggest that pigeons rely on experience gathered during the foraging session rather than on prior knowledge of patch quality or explicit cues, and they support the idea that foraging decisions emerge from continuous interaction with the local environment.

## Declarations

### Ethics approval

The experiments were carried out in compliance with the European Communities Council Directive of September, 22 2010 (2010/63/EU) and the specifications of the German law for the prevention of cruelty to animals and was approved by the animal ethics committee of the Landesamt für Natur, Umwelt und Verbraucherschutz (LANUV) NRW, Germany. We confirm that all methods were carried out in accordance with relevant guidelines and regulations and that the study was conducted in compliance with the ARRIVE guidelines.

### Code and Data availability

All data used in this study and the analysis scripts used for data processing and analysis are openly available on GitHub at [https://github.com/Guillermo-Hidalgo-Gadea/ForagingPigeonTracking]. Other software described in the methods section such as MotionPype, the PigeonSuperModel, Anipose and DeepLabCut is open source.

### Authors’ contributions

**Guillermo Hidalgo-Gadea:** Conceptualization, Methodology, Software, Formal analysis, Investigation, Data Curation, Writing - Original Draft, Writing - Review & Editing, Project administration. **Onur Güntürkün:** Resources, Writing - Review & Editing, Funding acquisition. **Mary Flaim:** Conceptualization, Methodology, Investigation, Resources, Writing - Review & Editing, Supervision. **Patrick Anselme:** Conceptualization, Methodology, Resources, Writing - Original Draft, Writing - Review & Editing, Supervision.

### Author Note

This study was supported by the Deutsche Forschungsgemeinschaft (DFG) under Grant Agreement GRK 274877981 (GHG), An1067/4-1 (PA), by the MERCATOR Research center at RUB under Grant Agreement No EX-2021-0018 (MF) as well as SFB 1280 project number 316803389 (OG) and the European Research Council (ERC-2020-ADG, grant agreement No. 101021354, AVIAN MIND) (OG). The authors have no conflicts of interest to disclose. We thank the members and students of the lab for their valuable support and feedback throughout this project. We are also grateful to the animal care staff for their dedicated work in maintaining the wellbeing of the pigeons.

