## Appendix A for "Continuous foraging behavior shapes patch-leaving decisions in pigeons: A 3D tracking study"

To accompany the Manuscript: Continuous foraging behavior shapes patch-leaving decisions in pigeons: A 3D tracking study

### Individual Performances

The following are individual performance reports of task engagement for every pigeon throughout the experiment, revealing interesting individual differences in foraging behavior across conditions. Out of the N = 12 pigeons tested, four tested consistently at ceiling level (P204, P430, P437, P600), three showed moderate foraging performances falling below criterion and recovering (P393, P434, P436). Two other pigeons started with high foraging performance but never reached the elevated platforms (P195, P428), and three other pigeons were excluded due to non-engagement with the experiment (P122, P589, P791). See the following sections for individual progress reports on these differences in engagement with the foraging setup.

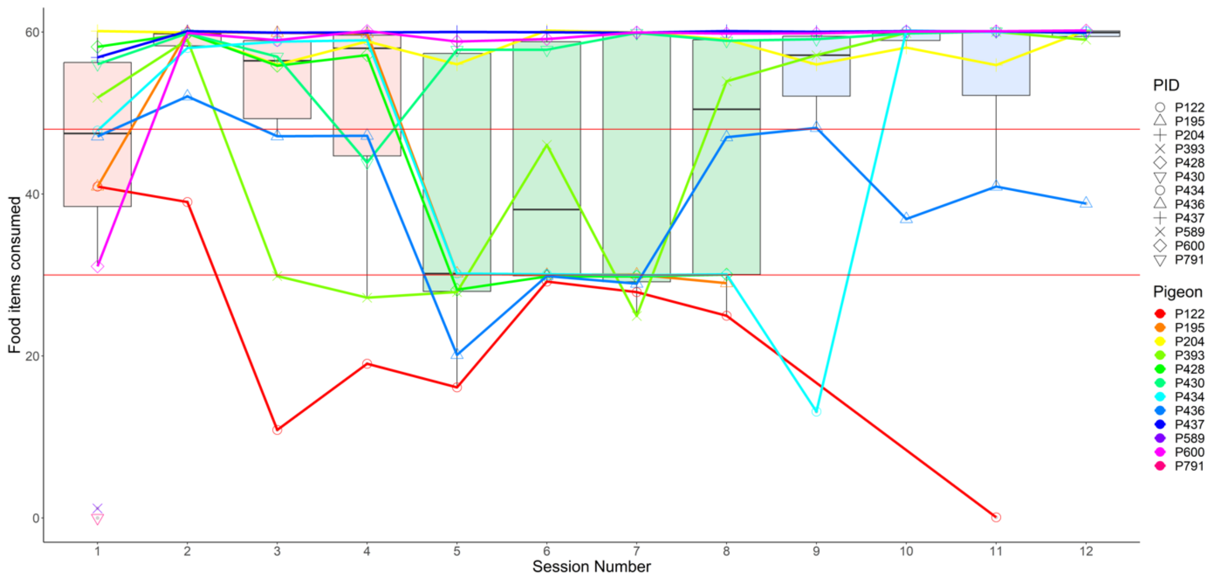


Figure A1: Foraging performance for each of the 12 pigeons throughout the 12 experimental sessions. Sessions 1-4 represent condition 0-0, with both platforms on the ground. Sessions 5-8 represent condition 0-75, with one platform elevated to 75cm. Sessions 9-12 represent condition 75-75, with both platforms elevated. Horizontal red lines indicate the 50% and 80% performance threshold out of 60 total food items. The boxplots represent the distribution of foraging performance per session.

**Pigeon P122**

P122 was a female pigeon, 2 years old at the start of the experiment, and with a normal body weight of 465 gram before food deprivation. It was housed individually throughout the experiment and tested during the second cohort. During training, P122 showed less interest towards the foraging boards compared to the other animals. Baiting the boards with sunflower seeds instead of green and yellow peas increased foraging activity during the first experimental sessions, but performance fell below 50% after the second session and the animal never visited the elevated platform. Visit latencies might not reflect foraging decisions due to the low task engagement. P122 should be excluded from the sample given the amount of missing data in unvisited platforms, the low foraging performance and different food items required (3g of pealed sunflower seeds: ~18kcal, 20% carbohydrate, 51% fat, 21% protein).

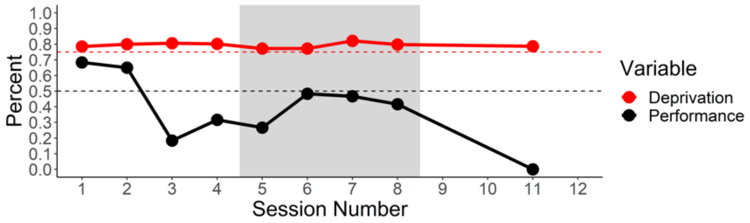


Figure A2. Individual task performance and food deprivation level for P122 throughout the experiment.

**Pigeon P791**

P791 was a male pigeon, 8 years old, and with a normal body weight of 520 gram before food deprivation. It was housed individually throughout the experiment and tested during the second cohort. During training, P791 showed no interest in the food items provided (green and yellow peas), and changing to sunflower seeds did not induce foraging behavior on the boards. During the first experimental session the pigeon showed normal exploratory behavior within the arena and frequently visited both ground platforms within the first 3 minutes, but the few visits and pecks on platform did not lead to any food item being consumed. P791 was exclude from further experimental sessions and returned to its regular feeding schedule.

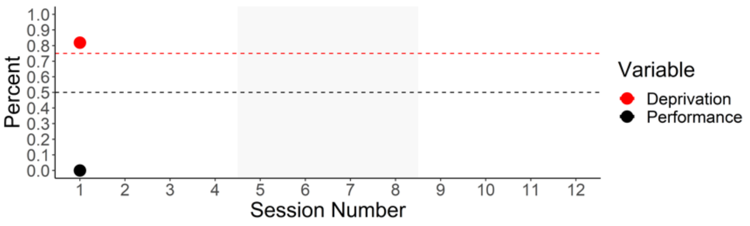


Figure A3. Individual task performance and food deprivation level for P791 throughout the experiment.

**Pigeon P589**

P589 was a male pigeon, 8 years old, and with a normal body weight of 490 gram before food deprivation. It was housed in aviary number 2 throughout the experiment and tested during the second cohort. During training, P589 showed no interest in the food items provided (green and yellow peas), and changing to sunflower seeds did not increase foraging performance. During the first experimental session the pigeon visited both ground platforms briefly within the first minute, but the few pecks on the platforms did not lead to any food item being consumed, and P589 starts exploring the arena without particular attention to the platforms after the first 2 minutes. P589 was exclude from further experimental sessions and returned to its regular feeding schedule.

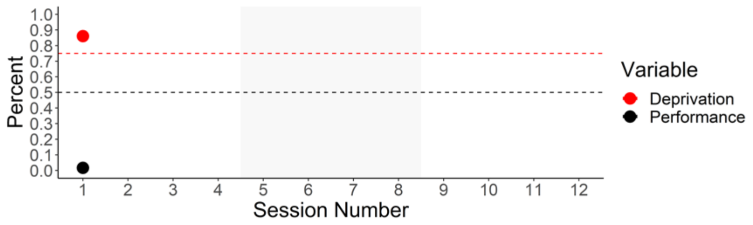


Figure A4. Individual task performance and food deprivation level for P589 throughout the experiment.

**Pigeon P195**

P195 was a female pigeon, 1 year old, and with a normal body weight of 480 gram before food deprivation. It was housed in aviary number 2 throughout the experiment and tested during the first cohort. After the first experimental session P195 fully depleted all ground patches, but during the 0-75 and the 75-75 conditions it never visited the elevated platforms. Note that even with optimal foraging performance during the first condition, by never visiting the elevated platform this animal contains missing data in later conditions and some data points are excluded from analyses regarding the elevated or even the second platform.

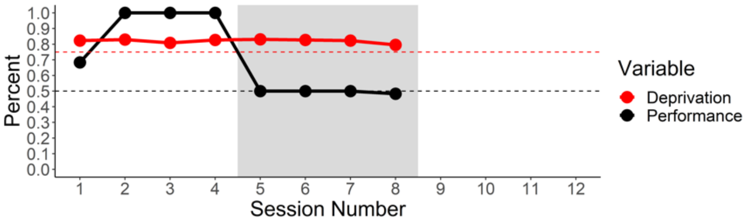


Figure A5. Individual task performance and food deprivation level for P195 throughout the experiment.

**Pigeon P204**

P204 was a female pigeon, 3 years old, and with a normal body weight of 460 gram before food deprivation. It was housed in aviary number 2 throughout the experiment and tested during the second cohort. Foraging performance of P204 was very high throughout the experiment, and the elevated platform was already visited and depleted during the first exposure.

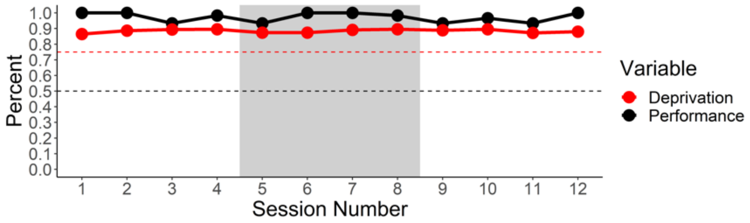


Figure A6. Individual task performance and food deprivation level for P204 throughout the experiment.

**Pigeon P393**

P393 was a female pigeon, 11 years old, and with a normal body weight of 400 gram before food deprivation. It was housed in aviary number 2 throughout the experiment and tested during the first cohort. Performance started high and fell after the second session, when it only depleted 50% of the food items while visiting both platforms. During the 0-75 condition, P393 visited and foraged from the elevated platform after the second exposure, but visited only the ground platform during session 7, with a second drop in performance without any big changes in food deprivation. During session 8 and after, it visited both platforms with a high foraging performance throughout the last condition. Note the missing data generated during sessions 5 and 7 by not visiting the second platform.

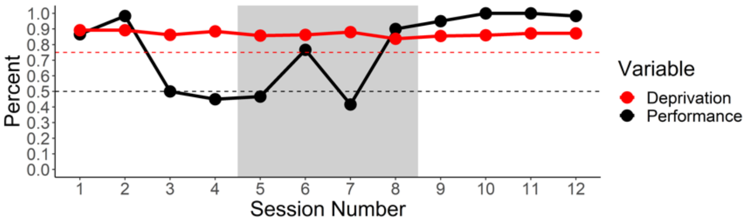


Figure A7. Individual task performance and food deprivation level for P393 throughout the experiment.

**Pigeon P428**

P428 was a female pigeon, 13 years old, and with a normal body weight of 400 gram before food deprivation. It was housed in aviary number 1 throughout the experiment and tested during the second cohort. This pigeon started at a high performance from the first session and almost depleted the ground platforms in every condition. By never visiting the elevated platform, foraging performance dropped to 50% during condition 0-75 and to 0% when both platforms were elevated. P428 did not visit the elevated platform after the 8th exposure and generated missing data from session 5 onwards for analyses regarding platform changes.

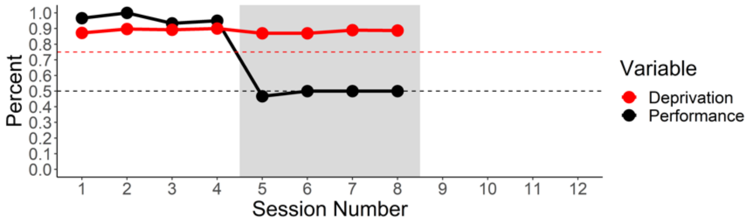


Figure A8. Individual task performance and food deprivation level for P428 throughout the experiment.

**Pigeon P430**

P430 was a female pigeon, 8 years old, and with a normal body weight of 456 gram before food deprivation. It was housed in aviary number 1 throughout the experiment and tested during the first cohort. P430 started with a high foraging performance from the first session, with a small dip during session 4, which corresponds to a small increase in food-deprived body weight, which might have affected foraging motivation. Performance recovered after that session and stayed very high throughout the rest of the experiment. P430 visited the elevated platform after the first exposure.

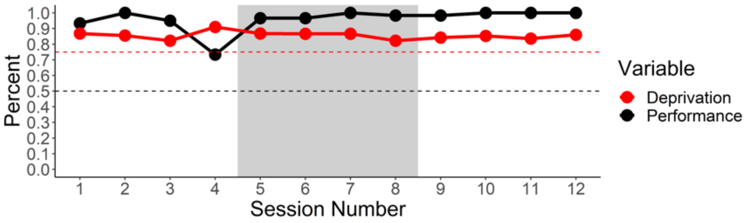


Figure A9. Individual task performance and food deprivation level for P430 throughout the experiment.

**Pigeon P434**

P434 was a female pigeon, 1 year old, and with a normal body weight of 524 gram before food deprivation. It was housed in aviary number 1 throughout the experiment and tested during the first cohort. Performance increased during the first sessions and dropped to 50% during the 0-75 condition, with the ground platform being perfectly depleted. When both platforms were elevated in session 9, P434 visited the first elevated platform with a latency of 18 minutes and only visited one of both platforms before the session ended 2 minutes after. From session 10 onwards, the animal perfectly depleted both platforms and latency to visit the elevated platform dropped to 6 seconds after the 5$^{th}$ exposure. Note the missing data generated during sessions 5 to 9 by only visiting one of both platforms. Note also that this animal was the largest in the sample and was highly food deprived throughout the experiment at an average of 77% of its normal body weight.

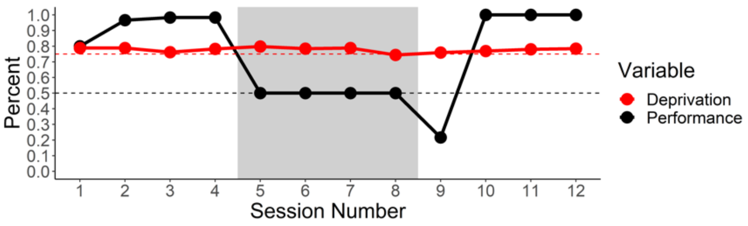


Figure A10. Individual task performance and food deprivation level for P434 throughout the experiment.

**Pigeon P436**

P436 was a female pigeon, 3 years old, and with a normal body weight of 415 gram before food deprivation. It was housed in aviary number 2 throughout the experiment and tested during the second cohort. Foraging performance was moderate for this animal without fully depleting the platforms. During condition 0-75, P436 visited the elevated platform during the 4$^{th}$ exposure with a moderate performance comparable to the first condition and fell below 75% for the remainder of the experiment while visiting and foraging from both platforms. Note the missing data generated during sessions 5, 6, and 7 by not visiting a second platform.

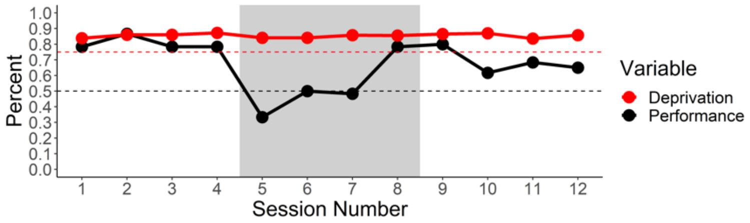


Figure A11. Individual task performance and food deprivation level for P436 throughout the experiment

**Pigeon P437**

P437 was a female pigeon, 1 year old, and with a normal body weight of 444 gram before food deprivation. It was housed in aviary number 2 throughout the experiment and tested during the first cohort. Performance was at ceiling level throughout the experiment, and P437 visited the elevated platform at first exposure.

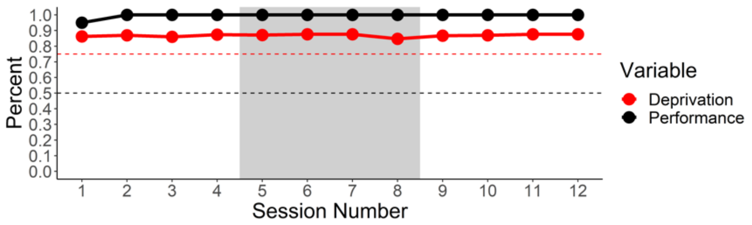


Figure A12. Individual task performance and food deprivation level for P437 throughout the experiment.

**Pigeon P600**

P600 was a female pigeon, 9 years old, and with a normal body weight of 485 gram before food deprivation. It was housed in aviary number 2 throughout the experiment and tested during the first cohort. Foraging performance started at 50% during the first session while visiting both platforms with a first latency of 17 seconds and a second platform latency of under 4 minutes, but performance reached and stayed at an average of 99% for the remaining sessions of the experiment. P600 visited the elevated platform at first exposure.

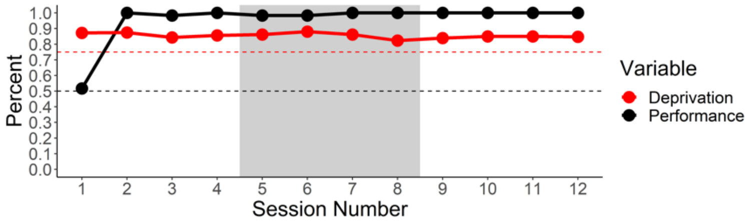


Figure A13. Individual task performance and food deprivation level for P600 throughout the experiment.
