## Appendix B for "Continuous foraging behavior shapes patch-leaving decisions in pigeons: A 3D tracking study"

To accompany the Manuscript: Continuous foraging behavior shapes patch-leaving decisions in pigeons: A 3D tracking study

### Model Coefficients

The following is a list of regression analyses used throughout the manuscript. Here we provide the complete regression equation and model coefficients. The data and scripts used for this analyses are openly available on GitHub at [https://github.com/Guillermo-Hidalgo-Gadea/ForagingPigeonTracking].

#### Model B1: Performance

Linear mixed effects regression on foraging performance by condition, with pigeons' ID as random intercept and session number, age, experimental weight, and free-feeding body weight as model covariates.

**R Output:**
```
Linear mixed model fit by REML. t-tests use Satterthwaite's method [
lmerModLmerTest]
Formula: food_consumed ~ condition + z_session_number_oncond + z_age +
    z_deprivation + z_normal_weight + (1 | PID)
   Data: data

REML criterion at convergence: 518.5

Scaled residuals:
    Min      1Q  Median      3Q     Max
-3.7458 -0.0163  0.1370  0.4824  1.7620

Random effects:
 Groups   Name        Variance Std.Dev.
 PID      (Intercept) 25.92    5.091
 Residual             39.14    6.256
Number of obs: 81, groups:  PID, 9

Fixed effects:
                        Estimate Std. Error      df t value Pr(>|t|)
(Intercept)              54.0521     2.0227  6.5295  26.723 6.46e-08 ***
condition0-75             3.1297     1.9331 72.3024   1.619   0.1098
condition75-75            3.2964     1.6849 72.1497   1.956   0.0543 .
z_session_number_oncond   0.1945     0.7011 67.4654   0.277   0.7823
z_age                     1.2431     1.8866  4.8276   0.659   0.5401
z_deprivation             1.2706     1.4334 52.2684   0.886   0.3795
z_normal_weight           1.7212     2.0207  6.0475   0.852   0.4268
---
Signif. codes:  0 '***' 0.001 '**' 0.01 '*' 0.05 '.' 0.1 ' ' 1
```

**Model B2: Transitions**

Linear mixed effects on the number of transitions between platforms by condition, with pigeons' ID as random intercept and session number, age, session performance, experimental weight, and free-feeding body weight as model covariates.

```
R Output:
Linear mixed model fit by REML. t-tests use Satterthwaite's method [
lmerModLmerTest]
Formula: num_transitions ~ condition + z_session_number_oncond + z_age +
    z_performance + z_deprivation + z_normal_weight + (1 | PID)
   Data: data

REML criterion at convergence: 595.3

Scaled residuals:
    Min      1Q  Median      3Q     Max
-2.1977 -0.4969 -0.0807  0.4283  5.1951

Random effects:
 Groups   Name        Variance Std.Dev.
 PID      (Intercept)   1.115   1.056
 Residual             131.640  11.473
Number of obs: 81, groups:  PID, 9

Fixed effects:
                         Estimate Std. Error        df t value Pr(>|t|)
(Intercept)              19.82252    2.01198  18.89366   9.852 7.06e-09 ***
condition0-75           -10.26165    3.41742  72.86605  -3.003  0.00366 **
condition75-75           -0.29859    2.98900  72.79210  -0.100  0.92070
z_session_number_oncond  -1.60326    1.28344  68.57893  -1.249  0.21584
z_age                    -4.19529    1.33833   3.38928  -3.135  0.04394 *
z_performance             0.66479    1.16240  49.24738   0.572  0.56999
z_deprivation             2.96414    1.77501  11.78160   1.670  0.12127
z_normal_weight           0.07124    1.70529   5.96226   0.042  0.96804
---
Signif. codes:  0 '***' 0.001 '**' 0.01 '*' 0.05 '.' 0.1 ' ' 1
```

**Model B3: Self-transitions**

Linear mixed effects regression on number of self-transitions by platform elevation, with pigeons' ID as random intercept and session number, age, session performance, experimental weight, and free-feeding body weight as model covariates.

**R Output:**

```
Linear mixed model fit by REML. t-tests use Satterthwaite's method [
lmerModLmerTest]
Formula: self_transitions_count ~ cond + z_session_number_oncond + z_age +
    z_performance + z_deprivation + z_normal_weight + (1 | PID)
   Data: data

REML criterion at convergence: 1035.6

Scaled residuals:
    Min      1Q  Median      3Q     Max
-2.3003 -0.6538 -0.1233  0.6425  3.9731

Random effects:
 Groups   Name        Variance Std.Dev.
 PID      (Intercept) 20.68    4.548
 Residual             36.31    6.026
Number of obs: 162, groups:  PID, 9

Fixed effects:
                         Estimate Std. Error         df t value Pr(>|t|)
(Intercept)              8.805890   1.712449   6.720134   5.142 0.001513 **
cond0-75 low            -0.816007   1.682158 151.064266  -0.485 0.628312
cond0-75 high            5.128438   1.682158 151.064266   3.049 0.002714 **
cond75-75 high           4.534306   1.186018 152.537447   3.823 0.000192 ***
z_session_number_oncond -0.009033   0.477802 147.125893  -0.019 0.984942
z_age                   -0.271682   1.633049   5.475828  -0.166 0.873841
z_performance            0.002697   0.499959 151.900343   0.005 0.995703
z_deprivation            1.340217   1.031607  99.248941   1.299 0.196901
z_normal_weight          0.581636   1.710088   6.600823   0.340 0.744331
---
Signif. codes:  0 '***' 0.001 '**' 0.01 '*' 0.05 '.' 0.1 ' ' 1
```

**Model B4: Peck-rate**

Linear mixed regression on peck rate by platform elevation, with pigeons' ID as random intercept and session number, age, session performance, experimental weight, and free-feeding body weight as model covariates. We control for the group-mean centered number of transitions within a session to remove between-condition differences and to ensure that the model controls only for within-condition effects.

**R Output:**

```
Linear mixed model fit by REML. t-tests use Satterthwaite's method [
lmerModLmerTest]
Formula:
total_peck_rate ~ cond + gmz_num_transitions + z_session_number_oncond +
    z_age + z_performance + z_deprivation + z_normal_weight +      (1 | PID)
   Data: data

REML criterion at convergence: 258.7

Scaled residuals:
    Min      1Q  Median      3Q     Max
-2.0538 -0.6550 -0.1938  0.4953  4.2116

Random effects:
 Groups   Name        Variance Std.Dev.
 PID      (Intercept) 0.01780  0.1334
 Residual             0.06895  0.2626
Number of obs: 1101, groups:  PID, 9

Fixed effects:
                          Estimate Std. Error         df t value Pr(>|t|)
(Intercept)              4.284e-01  4.665e-02  5.953e+00   9.184 9.82e-05 ***
cond0-75 low             9.947e-02  3.465e-02  1.091e+03   2.870  0.00418 **
cond0-75 high           -4.041e-03  3.790e-02  1.090e+03  -0.107  0.91510
cond75-75 high          -1.024e-01  1.884e-02  1.088e+03  -5.437 6.69e-08 ***
gmz_num_transitions     -6.889e-02  7.573e-03  1.088e+03  -9.097  < 2e-16 ***
z_session_number_oncond -1.131e-02  8.134e-03  1.088e+03  -1.391  0.16457
z_age                    1.983e-02  4.642e-02  5.776e+00   0.427  0.68466
z_performance            6.960e-03  1.096e-02  9.241e+02   0.635  0.52545
z_deprivation            4.905e-02  1.863e-02  5.951e+02   2.633  0.00868 **
z_normal_weight          9.481e-04  4.696e-02  6.243e+00   0.020  0.98452
---
Signif. codes:  0 '***' 0.001 '**' 0.01 '*' 0.05 '.' 0.1 ' ' 1
```

**Model B5: Inter-Peck Interval**

Linear mixed regression on the RMSSD variability of the inter-peck interval by platform elevation, with pigeons' ID as random intercept and session number, age, session performance, experimental weight, and free-feeding body weight as model covariates. We control for the group-mean centered number of transitions within a session to remove between-condition differences and to ensure that the model controls only within-condition effects.

**R Output:**

```
Linear mixed model fit by REML. t-tests use Satterthwaite's method [
lmerModLmerTest]
Formula:
total_IPI_rmssd ~ cond + gmz_num_transitions + z_session_number_oncond +
    z_age + z_performance + z_deprivation + z_normal_weight +      (1 | PID)
   Data: data

REML criterion at convergence: 4951.5

Scaled residuals:
    Min      1Q  Median      3Q     Max
-1.7336 -0.5480 -0.1969  0.2622 12.7513

Random effects:
 Groups   Name        Variance Std.Dev.
 PID      (Intercept) 0.09726  0.3119
 Residual             5.14774  2.2689
Number of obs: 1101, groups:  PID, 9

Fixed effects:
                          Estimate Std. Error         df t value Pr(>|t|)
(Intercept)                1.76241    0.14395    6.52920  12.243 9.59e-06 ***
cond0-75 low               0.02887    0.29554 1020.22430   0.098  0.92221
cond0-75 high              2.09230    0.32405 1050.68800   6.457 1.63e-10 ***
cond75-75 high            -0.24001    0.15795  691.89378  -1.520  0.12908
gmz_num_transitions       -0.19503    0.06505 1075.87530  -2.998  0.00278 **
z_session_number_oncond    0.02159    0.06988 1077.41055   0.309  0.75744
z_age                      0.05407    0.13399    4.62960   0.404  0.70449
z_performance             -0.23211    0.08424   93.42546  -2.755  0.00705 **
z_deprivation             -0.21410    0.12764   25.69473  -1.677  0.10560
z_normal_weight            0.08681    0.15131    7.04315   0.574  0.58402
---
Signif. codes:  0 '***' 0.001 '**' 0.01 '*' 0.05 '.' 0.1 ' ' 1
```

**Model B6: Inter-Peck Distance**

Linear mixed regression on the RMSSD variability of the inter-peck distances by platform elevation, with pigeons' ID as random intercept and session number, age, session performance, experimental weight, and free-feeding body weight as model covariates. We control for the group-mean centered number of transitions within a session to remove between-condition differences and to ensure that the model controls only within-condition effects.

**R Output:**

```
Linear mixed model fit by REML. t-tests use Satterthwaite's method [
lmerModLmerTest]
Formula:
total_IPD_rmssd ~ cond + gmz_num_transitions + z_session_number_oncond +
    z_age + z_performance + z_deprivation + z_normal_weight +      (1 | PID)
   Data: data

REML criterion at convergence: 12812.3

Scaled residuals:
    Min      1Q  Median      3Q     Max
-1.8132 -0.8083 -0.0897  0.5713  3.6623

Random effects:
 Groups   Name        Variance Std.Dev.
 PID      (Intercept)  142.2   11.92
 Residual             6928.6   83.24
Number of obs: 1101, groups:  PID, 9

Fixed effects:
                         Estimate Std. Error        df t value Pr(>|t|)
(Intercept)              107.7554     5.4036    9.1655  19.941 7.31e-09 ***
cond0-75 low              20.4350    10.8491 1043.6559   1.884   0.0599 .
cond0-75 high             20.0819    11.8941 1064.4556   1.688   0.0916 .
cond75-75 high           -30.7419     5.8031  793.0001  -5.298 1.52e-07 ***
gmz_num_transitions      -21.9601     2.3873 1082.0678  -9.199  < 2e-16 ***
z_session_number_oncond    0.1893     2.5645 1082.9468   0.074   0.9412
z_age                    -11.7703     5.0508    6.6372  -2.330   0.0545 .
z_performance             -0.9608     3.1086  137.9736  -0.309   0.7577
z_deprivation              3.9959     4.7359   38.8391   0.844   0.4040
z_normal_weight           -9.7636     5.6749    9.9459  -1.720   0.1162
---
Signif. codes:  0 '***' 0.001 '**' 0.01 '*' 0.05 '.' 0.1 ' ' 1
```

**Model B7: Area covered**

Linear mixed regression model on the log-transformed head-distance covered while foraging on a platform by the platform elevation, with pigeons' ID as random intercept and session number, age, session performance, experimental weight, and free-feeding body weight as model covariates. We control for the group-mean centered number of transitions within a session to remove between-condition differences and to ensure that the model controls only within-condition effects.

**R Output:**

```
Linear mixed model fit by REML. t-tests use Satterthwaite's method [
lmerModLmerTest]
Formula:
log(total_head_disp) ~ cond + gmz_num_transitions + z_session_number_oncond +
    z_age + z_performance + z_deprivation + z_normal_weight +      (1 | PID)
   Data: data

REML criterion at convergence: 2771.1

Scaled residuals:
    Min      1Q  Median      3Q     Max
-3.3971 -0.7146 -0.0401  0.7558  2.9757

Random effects:
 Groups   Name        Variance Std.Dev.
 PID      (Intercept) 0.06453  0.2540
 Residual             0.69318  0.8326
Number of obs: 1101, groups:  PID, 9

Fixed effects:
                          Estimate Std. Error         df t value Pr(>|t|)
(Intercept)              8.250e+00  9.341e-02  5.453e+00  88.324 8.12e-10 ***
cond0-75 low             3.310e-01  1.095e-01  1.086e+03   3.022  0.00257 **
cond0-75 high            1.253e+00  1.199e-01  1.089e+03  10.453  < 2e-16 ***
cond75-75 high           3.554e-01  5.931e-02  1.029e+03   5.992 2.86e-09 ***
gmz_num_transitions     -3.245e-01  2.398e-02  1.091e+03 -13.531  < 2e-16 ***
z_session_number_oncond -1.651e-02  2.576e-02  1.090e+03  -0.641  0.52176
z_age                    9.375e-03  9.181e-02  5.014e+00   0.102  0.92263
z_performance           -3.060e-02  3.381e-02  4.593e+02  -0.905  0.36592
z_deprivation           -8.855e-02  5.597e-02  1.593e+02  -1.582  0.11561
z_normal_weight          2.951e-02  9.549e-02  5.968e+00   0.309  0.76781
---
Signif. codes:  0 '***' 0.001 '**' 0.01 '*' 0.05 '.' 0.1 ' ' 1
```

**Model B8: Overpecking**

Linear mixed regression model on the overpecking ratio by the platform elevation, with pigeons' ID as random intercept and session number, age, session performance, experimental weight, and free-feeding body weight as model covariates. We control for the group-mean centered number of transitions within a session to remove between-condition differences and to ensure that the model controls only within-condition effects.

```
R Output:
Linear mixed model fit by REML. t-tests use Satterthwaite's method [
lmerModLmerTest]
Formula:
overpeck_ratio ~ cond + gmz_num_transitions + z_session_number_oncond +
    z_age + z_performance + z_deprivation + z_normal_weight +      (1 | PID)
   Data: data

REML criterion at convergence: -1657.4

Scaled residuals:
    Min      1Q  Median      3Q     Max
-3.1008 -0.6595 -0.0130  0.6424  4.5255

Random effects:
 Groups   Name        Variance Std.Dev.
 PID      (Intercept) 0.00000  0.0000
 Residual             0.01079  0.1039
Number of obs: 1026, groups:  PID, 9

Fixed effects:
                          Estimate Std. Error         df t value Pr(>|t|)
(Intercept)              1.917e-02  4.553e-03  1.016e+03   4.210 2.78e-05 ***
cond0-75 low            -4.110e-02  1.496e-02  1.016e+03  -2.748  0.00611 **
cond0-75 high            6.218e-02  1.472e-02  1.016e+03   4.224 2.62e-05 ***
cond75-75 high          -6.862e-03  7.216e-03  1.016e+03  -0.951  0.34190
gmz_num_transitions     -6.595e-03  3.092e-03  1.016e+03  -2.133  0.03314 *
z_session_number_oncond -1.192e-03  3.289e-03  1.016e+03  -0.362  0.71712
z_age                   -8.800e-03  3.669e-03  1.016e+03  -2.399  0.01663 *
z_performance            3.091e-03  3.402e-03  1.016e+03   0.909  0.36374
z_deprivation           -1.835e-04  4.494e-03  1.016e+03  -0.041  0.96743
z_normal_weight         -2.509e-03  4.784e-03  1.016e+03  -0.525  0.60003
---
Signif. codes:  0 '***' 0.001 '**' 0.01 '*' 0.05 '.' 0.1 ' ' 1
```
